# A conserved role of the duplicated *Masculinizer* gene in sex determination of the Mediterranean flour moth, *Ephestia kuehniella*

**DOI:** 10.1101/2021.02.15.431251

**Authors:** Sander Visser, Anna Voleníková, Petr Nguyen, Eveline C. Verhulst, František Marec

## Abstract

Sex determination in the silkworm, *Bombyx mori*, is based on *Feminizer* (*Fem*), a W-linked *Fem* piRNA that triggers female development in WZ individuals, and the Z-linked *Masculinizer* (*Masc*), which initiates male development and dosage compensation in ZZ individuals. While *Fem* piRNA is missing in a close relative of *B. mori*, *Masc* determines sex in several representatives of distant lepidopteran lineages. We studied the molecular mechanisms of sex determination in the Mediterranean flour moth, *Ephestia kuehniella* (Pyralidae). We identified an *E. kuehniella Masc* ortholog, *EkMasc*, and its paralog resulting from a recent duplication, *EkMascB*. Both genes are located on the Z chromosome and encode a similar Masc protein that contains two conserved domains but has lost the conserved double zinc finger domain. We developed PCR-based genetic sexing and demonstrated a peak in the expression of *EkMasc* and *EkMascB* genes only in early male embryos. Simultaneous knock-down experiments of both *EkMasc* and *EkMascB* using RNAi during early embryogenesis led to a shift from male- to female-specific splicing of the *E. kuehniella doublese*x gene (*Ekdsx*), their downstream effector, in ZZ embryos and resulted in a strong female-biased sex-ratio. Our results thus confirmed the conserved role of both *EkMasc* and *EkMascB* genes in masculinization. We suggest that the C-terminal proline-rich domain, we have identified in all functionally confirmed Masc proteins, in conjunction with the masculinizing domain, is important for transcriptional regulation of sex determination in Lepidoptera. The function of the Masc double zinc finger domain is still unknown, but appears to have been lost in *E. kuehniella*.

**Author summary:** The sex-determining cascade in the silkworm, *Bombyx mori*, differs greatly from those of other insects. In *B. mori*, female development is initiated by *Fem* piRNA expressed from the W chromosome during early embryogenesis. *Fem* piRNA silences *Masculinizer* (*Masc*) thereby blocking the male pathway resulting in female development. It is currently unknown whether this cascade is conserved across Lepidoptera. In the Mediterranean flour moth, *Ephestia kuehniella*, we identified an ortholog of *Masc* and discovered its functional duplication on the Z chromosome, which has not yet been found in any other lepidopteran species. We provide two lines of evidence that both the *EkMasc* and *EkMascB* genes play an essential role in masculinization: (i) they show a peak of expression during early embryogenesis in ZZ but not in WZ embryos and (ii) their silencing by RNAi results in female-specific splicing of the *E. kuehniella doublesex* gene (*Ekdsx*) in ZZ embryos and in a female-biased sex ratio. Our results suggest a conserved role of the duplicated *Masc* gene in sex determination of *E. kuehniella*.

## Introduction

Sex determination in insects is under the control of a cascade of genes, each affecting the expression or splicing of the next gene in the pathway [1]. This cascade evolved from the bottom up with the most conserved gene, *doublesex* (*dsx*), at the bottom of the cascade [2] present in all insects studied to date [3–5]. The *dsx* gene is sex-specifically spliced by an upstream splicing factor, e.g. *transformer* (*tra*) in Hymenoptera, Coleoptera, and derived Brachycera (a suborder of Diptera) [6, 7]. In turn, the activity of *tra* is affected by the presence of a primary signal gene which can either activate or inactivate *tra* thereby acting as the initiator of sex determination. The upstream sex determination cascade seems to undergo rapid evolutionary change, as these genes are often replaced, duplicated and reshuffled, as have been shown in multiple species. For example, in Hymenoptera, the feminizing gene *wasp overruler of masculinization* (*wom*) in the jewel wasp, *Nasonia vitripennis*, is a novel gene originating from a P53 gene duplication, but has also been very recently duplicated, with both copies of the gene thought to be functional [8]. The gene initiating sexual differentiation in the honeybee *Apis mellifera*, *complementary sex determiner* (*csd*), originated via a duplication of the *feminizer* (*fem*) gene, an ortholog of *tra* [9, 10]. In addition, duplications of *tra*/*fem* have been detected in many other hymenopteran species, though their role in sex determination is unknown [6, 11, 12]. Also in the housefly *Musca domestica* (Diptera), the masculinizing gene *male determiner* (*Mdmd*) originated through a gene duplication and is located in a locus containing multiple pseudocopies of the gene [13].

The insect order Lepidoptera (moths and butterflies) includes pollinators and several other beneficial species of high economic importance, e.g. the silkworm *Bombyx mori*, but also a large number of major pests of agricultural crops, such as the diamondback moth *Plutella xylostella* [14]. The lepidopteran sex determination cascade has diverged from other holometabolous insects as *tra* was presumably lost in this order [6, 15]. The model species in lepidopteran sex determination research is *B. mori*, which has a WZ/ZZ sex chromosome constitution with a dominant feminizing W chromosome (Hasimoto 1933 as cited in [16]). In ZZ individuals, the Z-linked masculinizing gene *Masculinizer* (*Masc*) initiates male development and dosage compensation. In WZ *B. mori* individuals, the W-linked *Feminizer* (*Fem*) piRNA targets the *Masc* mRNA resulting in its degradation [17]. In the absence of Masc protein, the *B. mori doublesex* (*Bmdsx*) gene undergoes female-specific splicing resulting in female development. In the absence of the W chromosome, *Masc* is not suppressed by *Fem* piRNA and promotes male-specific splicing of *Bmdsx*, although the pathway through which Masc affects splicing of *Bmdsx* is currently unknown. The Masc protein contains two zinc finger motifs in the N-terminus [17], a bipartite nuclear localization signal (bNLS) [18], and a masculinizing region [19]. An additional segment located in the C-terminus of the protein is essential for masculinization and dosage compensation but the exact functional motif has yet to be determined [19, 20]. Masculinization through *Masc* seems to be a conserved feature of the lepidopteran sex determination mechanism, and has been confirmed in *Trilocha varians* (Bombycidae) [21], *Ostrinia furnacalis* (Crambidae) [22, 23], *Agrotis ipsilon* (Noctuidae) [24], and *P. xylostella* (Plutellidae) [25]. In addition, regulation of dosage compensation has been confirmed in *O. furnacalis* [22] and suggested for *T. varians* [26] and *P. xylostella* [25]. Contrary to masculinization by *Masc*, feminization through *Fem* piRNA is not conserved in *T. varians*, a close relative of *B. mori* [21].

Considering the economic importance of Lepidoptera, extensive research into sex determination mechanisms is still lacking in this large insect order. Therefore, we set out to identify the sex determination mechanism in the Mediterranean flour moth, *Ephestia kuehniella*. This moth was one of early models of genetics in Lepidoptera (see *Anagasta kühniella* in [27]) and later a model for sex chromosome research [28]. It used to be a serious pest of dried food products, in particular flour, but due to improved hygiene regulations, it is no longer considered a major pest [29]. Nowadays, *E. kuehniella* is widely accepted as a factitious host that is easy to rear in large quantities, and thus its eggs and larvae are used as a food source for a broad range of beneficial arthropods used in biological control [30–32]. Abiotic- and biotic factors affecting the growth and development of *E. kuehniella* have been studied extensively [33, 34], but egg production could be considerably increased by alteration of the sex ratio of the population. Males are able to sire the complete offspring of at least 9 females [35], and therefore the number of males in the population can be reduced while retaining complete fertilization of the population. Therefore, identification of the sex determining mechanism in *E. kuehniella* may potentially provide candidate target genes to alter the sex ratio, thereby increasing egg production.

Here, we identified a *Masc* ortholog in *E. kuehniella*, *EkMasc*, and its paralog resulting from a recent duplication, *EkMascB*. We developed a PCR-based genetic sexing method in *E. kuehniella* to be able to analyze *EkMasc* and *EkMascB* function using RNAi during early embryogenesis. Our results show that despite the loss of the zinc finger motifs, their role in masculinization is intact. In addition, we characterized a male-specific splice variant of *EkMasc* and *EkMascB*, skipping the exon containing the masculinizing region. Finally, we identified conserved regions in the C-terminus of all functionally confirmed lepidopteran Masc proteins that are likely involved in masculinization. Overall, our results substantially add to the understanding of the core genes of the lepidopteran sex determination cascade and its evolution, but also provide target genes for sex ratio optimization for mass rearing purposes.

## Results

### Identification of *EkMasc* and its paralog *EkMascB*

Initial sequences of *Masculinizer* (*EkMasc*) in *E. kuehniella* were identified using reverse transcription PCR (RT-PCR). Full cDNA sequences were subsequently obtained using 5’ and 3’ rapid amplification of cDNA ends (RACE) followed by sequencing. Two variants of *EkMasc* were consistently found, but we were not sure if they were allelic variants or duplication of the *Masc* gene. A BLASTn search of the two complete cDNA sequences against the draft genome assembly of *E. kuehniella* revealed a single scaffold with both copies of the gene, strongly suggesting that the two sequences obtained are not allelic variants but signatures of a real *EkMasc* duplication, which we denoted *EkMascB*. The open reading frames of *EkMasc* and *EkMascB* are in opposite orientation (tail-tail) with ∼23 kb distance between the two genes (Fig 1A). In addition, an ortholog of the *6-phosphogluconate dehydrogenase* (*Ek6-Pgd*) gene was identified in the same scaffold in close proximity to *EkMasc*. Within the intron of *Ek6-Pgd* between exons 7 and 8, a complete mariner transposon lacking an intact open reading frame was identified and a similar but partial mariner transposon was identified between the *EkMasc* and *EkMascB* genes, containing only the 3’ region of the transposon. The two mariner transposon sequences share 93% sequence identity.

**Fig 1.**
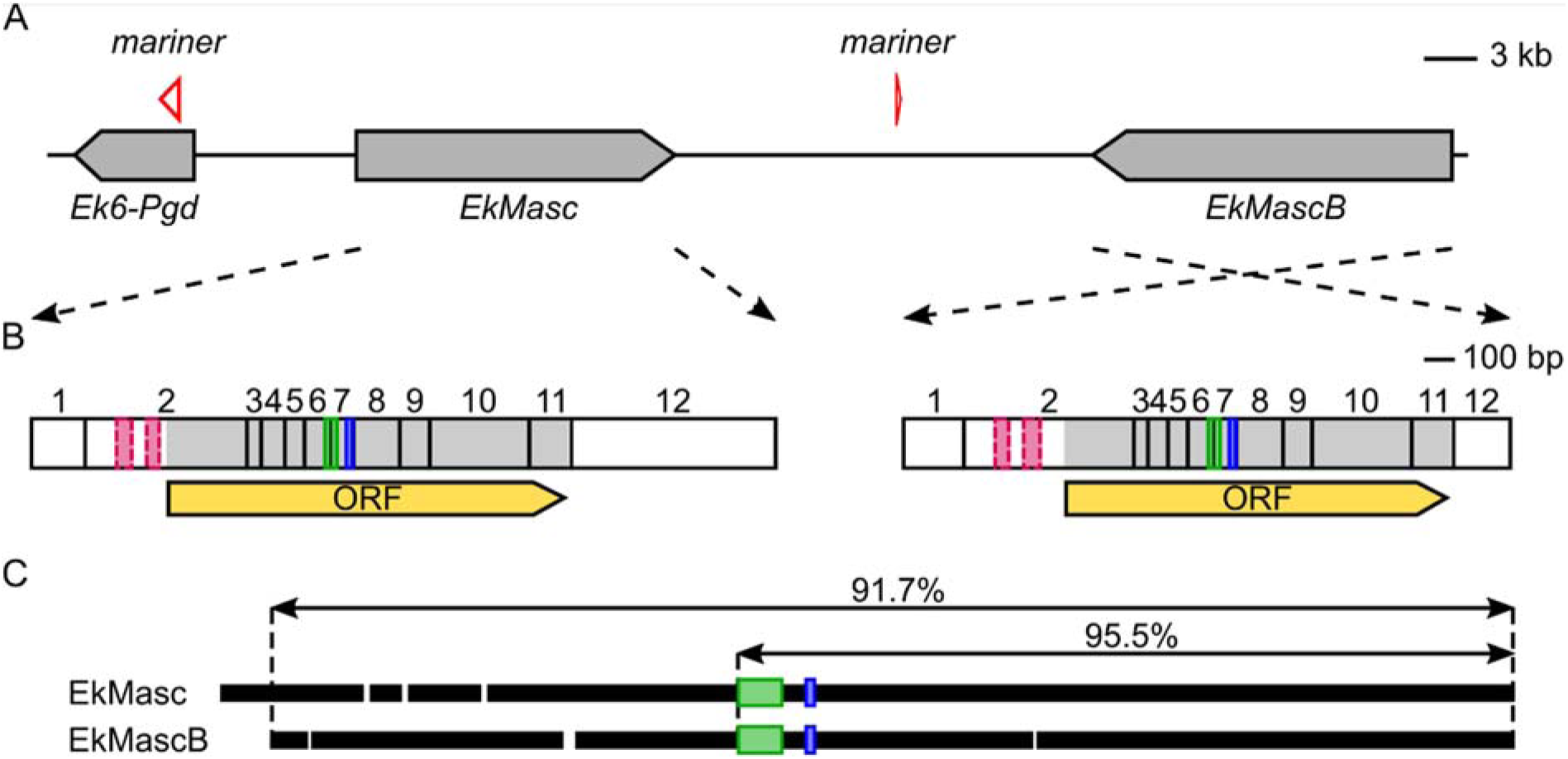
Genomic organization of EkMasc and EkMascB genes in Ephestia kuehniella. (**A**) Graphical representation of *EkMasc* and *EkMascB* on the assembled scaffold with an ortholog of the *6-Pgd* gene (*Ek6-Pgd*) upstream of *EkMasc* and two inactive mariner transposase genes. (**B**) Representation of complete transcripts of *EkMasc* and *EkMascB* with the exons, open reading frame (ORF; yellow), bipartite nuclear localization signal (bNLS; green), masculinizing domain (blue), and degenerated zinc finger domains in exon 2 (dashed pink) indicated. (**C**) Alignment of the EkMasc and EkMascB protein sequences showing the amount of homology between the two proteins, white spaces are indels, green and blue boxes represent the bNLS and masculinizing domains, respectively.

### Localization of *EkMasc* and *EkMascB* on the Z chromosome

In *B. mori* and *Plutella xylostella*, *Masc* is located on the Z chromosome [17, 25]. As with most lepidopterans, the Z chromosome in *E. kuehniella* is present in one copy in females but in two copies in males [36]. Gene content of the Z chromosome between lepidopteran species is generally highly conserved [37–39], but no information regarding the content of the Z chromosome of *E. kuehniella* is available. Therefore, we verified the location of *EkMasc* on the Z chromosome in *E. kuehniella* using quantitative real-time PCR (qPCR) on genomic DNA. We tested two hypotheses regarding the location of the two genes, (i) *EkMasc* and *EkMascB* are Z-linked resulting in a female to male ratio of 0.5 and (ii) both genes are autosomal resulting in a ratio of 1 [37]. Data from qPCR with genomic DNA of *E. kuehniella* females and males targeting both *EkMasc* and *EkMascB* genes at the same time were normalized against the autosomal reference gene, *Acetylcholinesterase 2* (*Ace-2*) (S1 Fig). An unpaired two-tailed *t*- test for unequal variances showed a statistically significant difference in the female and male normalized quantities (*P* = 0.0049), thus ruling out the autosomal hypothesis. In addition, a comparison of the male normalized quantities against the female normalized quantities multiplied by 2 showed no significant difference between ratios (*P* = 0.8415), therefore strongly suggesting that both *EkMasc* and *EkMascB* are located on the Z chromosome. These normalized *EkMasc*+*EkMascB* quantities were compared between females and males showing that the normalized female to male ratio of *EkMasc*+*EkMascB* in *E. kuehniella* was 0.505 ± 0.032.

Our Southern hybridization data consistently showed that the two signals present in both sexes are stronger in males than in females (S2 Fig), which is in line with the qPCR results and confirms the Z-linkage of *EkMasc* and *EkMascB*. This further corroborates our conclusion that there are two *EkMasc* copies, because the single Z chromosome in females shows two bands in Southern hybridization, ruling out allelic variation.

### Nucleotide and protein comparison of *EkMasc* and *EkMascB*

*EkMasc* and *EkMascB* complete cDNA sequences were used in a BLASTn search against the draft genome to identify and annotate the exons of both genes. Next, we aligned *EkMasc* to *EkMascB* to compare the structure of both genes. All exons of *EkMasc* were also present in *EkMascB*, and both genes contained an open reading frame (S3 Fig). Divergence between the two genes, i.e. single nucleotide polymorphisms (SNPs), insertions and deletions (indels), was strongest in the 5’- and 3’- UTRs and in the intron sequences. We obtained five splice variants for *EkMasc* and seven splice variants for *EkMascB* from sequencing cloned RT-PCR and RACE-PCR products. Six of these splice variants (two for *EkMasc* and four for *EkMascB*) appeared to be present at a very low frequency as they were not visibly amplified and contained early stop codons as a consequence of exon-skipping, intron-retention, or premature polyadenylation. Therefore, we considered these six variants biologically non-relevant.

The remaining six splice variants (three for each gene) were abundantly present. In both genes, the first and second splice variant, *EkMasc-1* and *EkMasc-2* (and *EkMascB-1* and *EkMascB-2*), contained the bipartite nuclear localization signal (bNLS) [18] and two cysteine residues that are presumably essential for masculinization [19] and differ only in splicing of the terminal (non-coding) exon (Fig 1B; S3 Fig). These two splice variants are the only variants detected during early embryogenesis (Fig 2A). The third splice variant, termed masculinizing domain skipping *Masc* (*EkMasc^ms^* and *EkMascB^ms^*) skips the exon containing the masculinizing domain (exon VII), resulting in an early stop codon (S3 Fig). This splice variant appears later during development and exclusively in males (Fig 2A). Splice forms containing the masculinizing domain appear to predominate in the testes, while *EkMasc^ms^* and *EkMascB^ms^* appear to predominate in somatic tissues (Fig 2B). Due to the importance of the masculinizing domain for Masc function, we investigated whether the *Masc^ms^* splice form is present in other lepidopterans or if it is specific to *E. kuehniella*. The presence and distribution of the *Masc^ms^* splice form was tested in the closely related Indian meal moth, *Plodia interpunctella*, and the distant codling moth, *Cydia pomonella*. In both species, *Masc^ms^* is found in male somatic tissues but not in the testes, and results in a premature stop codon, similar to *E. kuehniella* (Fig 2C, D). In *P. interpunctella*, we also found that the exon containing the masculinizing region is shorter than the annotated *PiMasc* sequence, resulting in an exon of similar size to the homologous exon in *EkMasc*.

**Fig 2.**
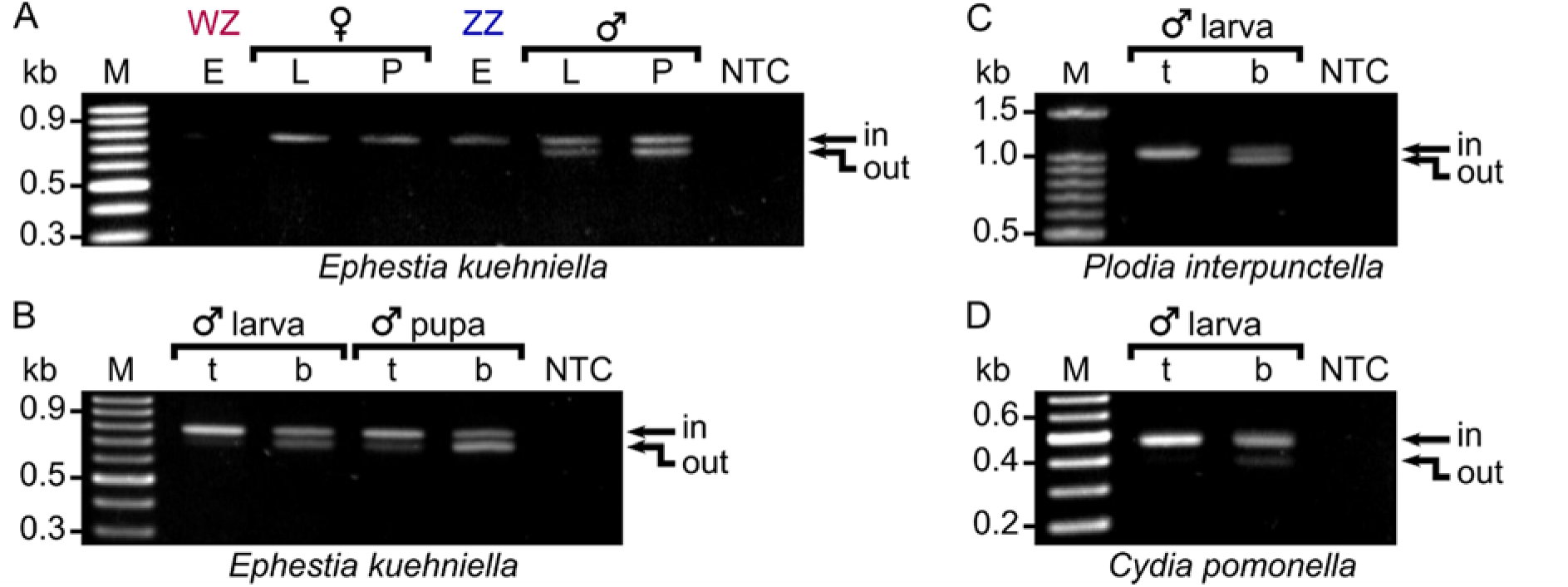
Alternative splicing of *Masc* in different life stages of *Ephestia kuehniella* and different tissues in *E.kuehniella*, *Plodia interpunctella*, and *Cydia pomonella*. (**A**) Alternative splicing of *EkMasc* and *EkMascB* in WZ and ZZ samples of different life stages, embryo 16 hpo (E), larva (L), and pupa (P), where the exon containing the masculinization domain (exon VII) is either spliced in or spliced out. (**B**) Male testis (t) and whole body samples minus the testis (b) show different splicing patterns in *E. kuehniella* with the testis sample showing predominantly splicing-in of exon VII in both larva and pupa stage, and the body samples showing both splice types. This splice pattern is conserved in males of the closely related *P. interpunctella* (**C**) and the more distant *C. pomonella* (**D**).

The complete *EkMasc* mRNA sequence containing the full open reading frame is 2425 bp (*EkMasc-1*) or 2667 bp long (*EkMasc-2*) depending on the splice variation in the 3’ terminal exon of the gene and encodes a 436 amino acid protein. Similarly, the complete mRNA sequence of *EkMascB* is 2346 bp (*EkMascB-1*) or 2644 bp (*EkMascB-2*) long and encodes a slightly shorter protein of 419 amino acids. Comparison of EkMasc with EkMascB shows that they share 91.7% homology over the entire length of the proteins and 95.5% homology in the region likely to be important for masculinization [19], as seen in Fig 1C. EkMasc and EkMascB differ from each other by 6 indels and 23 substitutions (S4 Fig). Both the male determining domain [19] and the bNLS domain [18] are present in both EkMasc and EkMascB. In addition, two substitutions are present within the spacer sequence of the bNLS domain, but the domain itself and the masculinizing domain share complete homology. Interestingly, the two zinc-finger motifs present in the N-terminus of BmMasc [17] and most other lepidopteran Masc proteins [21, 22, 24, 25] were lost in both EkMasc and EkMascB. The residues of the double zinc finger motif remain part of the transcripts of both genes, yet the sequences have degenerated independently, and the accumulation of mutations has shifted the open reading frame of both genes downstream of the degenerated domains (Fig 1B).

### Functional analysis of *EkMasc* and *EkMascB*

Expression levels of all *EkMasc* and *EkMascB* splice forms were measured during embryogenesis using quantitative reverse transcription PCR (qRT-PCR) to determine sex-specific expression patterns for both genes separately. Embryos were sexed using an in-house developed PCR-based sexing system that shows an additional band in female samples (S5 Fig).

Both *EkMasc* and *EkMascB* were expressed throughout embryogenesis and both genes followed the same pattern of expression level differences between the sexes (Fig 3). At 12 hours post oviposition (hpo), no significant difference in expression level of both genes was observed between the sexes. After 12 hpo, expression levels of *EkMasc* and *EkMascB* increased in males until they reached a maximum around 16 hpo, while the expression levels decreased in females simultaneously. After 16 hpo, expression levels decreased in males and reached levels comparable to females at approximately 24 hpo. We performed an unpaired two-tailed *t*-test with unequal variances to test for differences between the sexes, showing statistically significant differences at 14–22 hpo for both genes (S1 Table).

**Fig 3.**
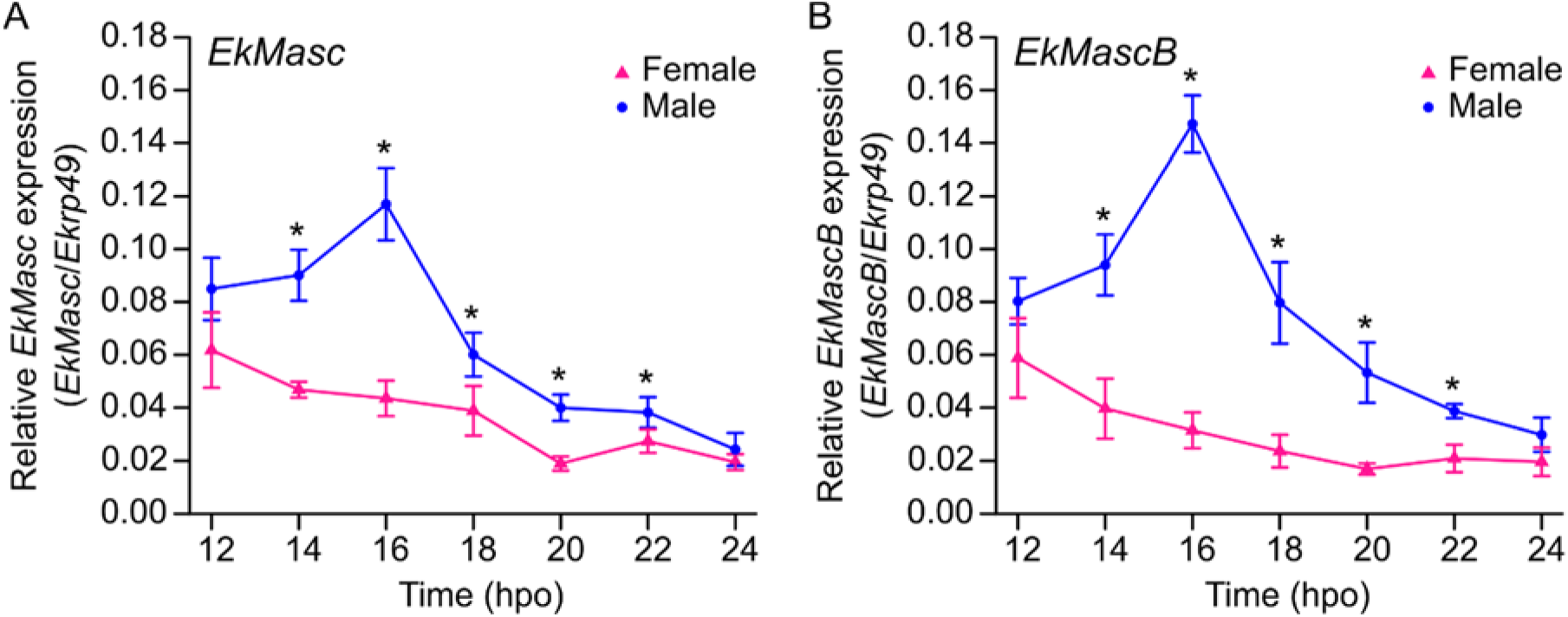
Expression of *EkMasc* (A) and *EkMascB* (B) during early embryogenesis in female (pink) and male (blue) samples of *Ephestia kuehniella*. Each time point has at least 3 biological replicates. Significant differences between sexes are indicated (*; unpaired two-tailed *t*-test; *P* < 0.05) and errors bars indicate standard deviation.

To verify the function of *EkMasc* and *EkMascB*, we performed a simultaneous knock-down of both genes by injecting short interfering RNAs (siRNAs) targeting a conserved region into eggs 1–2 hpo. We performed two separate knock-down experiments using siRNA targeting either exon II (siMasc_II) or exon VII (siMasc_VII), which contains the masculinizing domain (S6 Fig). For the third treatment, we injected a negative control siRNA designed against the *green fluorescent protein* (*GFP*) gene, as previously done [17]. We assessed knock-down efficiency by qRT-PCR at 16 hpo (Fig 4; S2 Table), the peak of *EkMasc* and *EkMascB* expression levels, as this is presumably the time point at which the expression of *EkMasc* and *EkMascB* is essential for male development. The average expression level of *EkMasc* in ZZ embryos after injection with siMasc_II and siMasc_VII was reduced to 51.6% (*P* = 0.0311) and 52.0% (*P* = 0.0300), respectively (Fig 4A), and *EkMascB* expression in ZZ embryos was reduced to 41.8% (*P* = 0.0137) and 39.1% (*P* = 0.0164), respectively (Fig 4B). In WZ embryos, expression of *EkMasc* was reduced to 65.4% (*P* = 0.0335) and 50.7% (*P* = 0.0034) after injection with siMasc_II and siMasc_VII, respectively, while *EkMascB* showed reduced expression levels of 62.8% (*P* = 0.0416) and 49.5% (*P* = 0.0090), respectively (Fig 4C, D).

**Fig 4.**
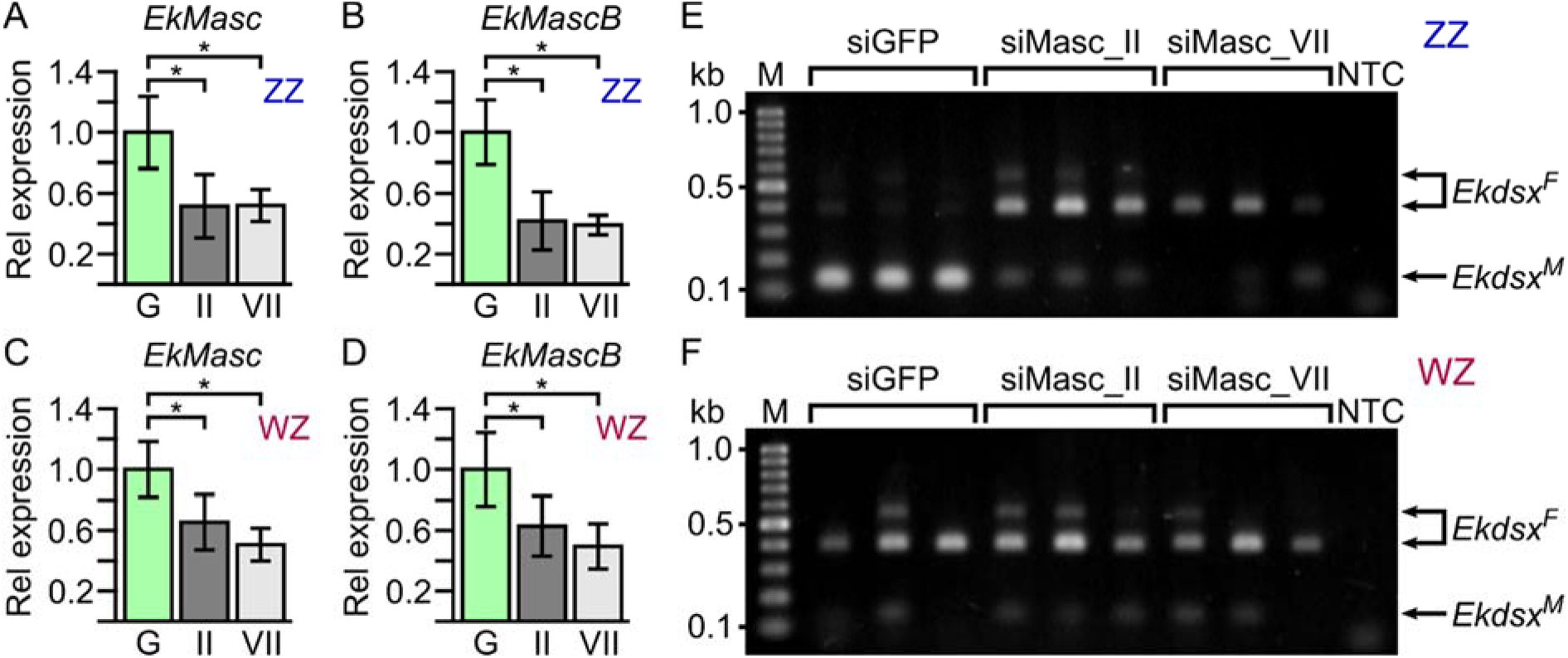
RNAi knock-down effects in ZZ and WZ *Ephestia kuehniella* individuals 16 hours post oviposition. The fold-change in expression levels of *EkMasc* (**A** and **C**) and *EkMascB* (**B** and **D**) in ZZ (**A** and **B**) and WZ (**C** and **D**) individuals injected with siMasc_II (si_II) or siMasc_VII (si_VII) are shown relative to the control injected (siGFP) individuals. Statistically significant differences are indicated by * (unpaired one-tailed *t*-test; *P* < 0.05; S2 Table), error bars indicate standard deviation. In **E** and **F**, effects of sex-specific splicing of *Ekdsx* 48 hpo after injection with control, siMasc_II, and siMasc_VII are shown in ZZ and WZ individuals, respectively. Note that splicing of *Ekdsx* shifts from predominantly male-specific in the control to mostly female-specific in siMasc_II- and siMasc_VII-treated ZZ individuals (**E**), while splicing does not differ between treatments in WZ individuals (**F**).

The *doublesex* (*dsx*) gene is often used as indicator of sexual development in knock-down experiments involving sex determination genes [21, 25]. We identified an ortholog of this gene in *E. kuehniella*, *Ekdsx*, and confirmed its sex-specific splicing using RT-PCR in pupae (S7 Fig). A single male-specific splice form was identified in males and two dominant female-specific splice forms were identified in females (S7A, B Fig). This is similar to the splicing structure identified in *B. mori* [40, 41]. In addition, a third female-specific splice form was observed in females (S7C Fig), but we failed to sequence this third female-specific splice form, and thus its splicing structure is currently unknown. Splicing of *Ekdsx* was female-specific in all early embryos, but transitions to male-specific splicing were observed only in ZZ individuals, 16–18 hpo at 21–22°C (S7D Fig). After injection of siRNA, sex-specific splicing of *Ekdsx* was assessed at 48 hpo to ensure that *Ekdsx* splicing stabilized in either the female- or male-specific isoform(s). WZ individuals injected with any of the three siRNAs (control siGFP, siMasc_II, and siMasc_VII) showed predominantly female-specific splicing of *Ekdsx* as expected (Fig 4F). ZZ individuals injected with control siGFP showed predominantly male-specific splicing of *Ekdsx* while those injected with siMasc_II or siMasc_VII showed predominantly female-specific splicing of *Ekdsx* (Fig 4E) comparable to WZ individuals. Approximately 400 eggs injected with siGFP were left to develop of which 12% (48/400) hatched and survived to adulthood. Similarly, approximately 10% (49/500) of the eggs injected with siMasc_II and approximately 8% (48/620) of the eggs injected with siMasc_VII hatched and survived to adulthood. Adults were phenotypically sexed based on external genitalia, revealing sex-ratios not significantly differing from 0.5 in those injected with siGFP (*P* = 0.5637) and siMasc_II (*P* = 0.8864) (Fig 5). Adults developing from embryos injected with siMasc_VII showed a strong female-biased sex-ratio of 0.85, which differed significantly from 0.5 (*P* = 9.226e-07). The heads of females that developed from siMasc_VII-injected embryos were used to test the genetic sex of the individual, which showed that all phenotypic females have WZ sex chromosomes, suggesting a male-killing effect of RNAi with siMasc_VII.

**Fig 5.**
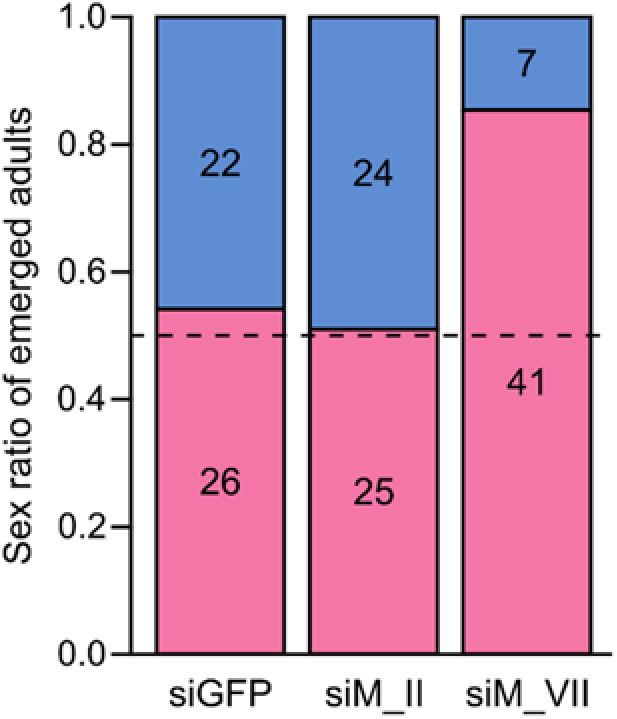
Sex ratio of emerged *Ephestia kuehniella* adults injected as 1–2 hpo embryos with siGFP, siMasc_II (siM_II), and siMasc_VII (siM_VII). The number of adult females (pink) and males (blue) emerged in each treatment are indicated in the figure. The sex ratio (f : m) of emerged adults injected with siGFP or siMasc_II is approximately 0.5 (indicated by a dashed line), while the sex ratio of siMasc_VII-injected individuals is strongly female-biased (> 0.8).

### Masculinizer comparison in Lepidoptera

We compared all published and functionally confirmed Masc protein sequences to identify any additional conserved domains and/or features of the proteins. Previously, three conserved domains were identified in BmMasc: (1) the double zinc finger domain [17], (2) the bNLS domain [18], and (3) the masculinizing domain [19]. The two cysteine residues that define the masculinizing domain and the bNLS domain are present in all species. The double zinc finger motif is conserved in most species with the exceptions of *E. kuehniella*, which lost both motifs in both copies of the gene, and potentially in *Trilocha varians* that has a deletion in the second zinc finger [21].

After aligning all functionally confirmed lepidopteran Masc protein sequences, we observed a relatively high level of proline residues in all proteins (S8 Fig). The average proline content of lepidopteran proteins is 5.52% (*B. mori*), 5.86% (*P. xylostella*), and 5.35% (*P. interpunctella*), while the proline content in their respective Masc proteins is 11.56%, 17.37%, and 13.73%. Given the less than 0.4% variation in the proline contribution across the total protein databases between the species, we assume that *E. kuehniella* has similar proline levels across all proteins, whereas EkMasc and EkMascB consist of 16.09% and 17.22% proline residues, respectively. To determine the distribution of the proline residues across the protein sequence, we performed a sliding window analysis with window size 25 on the Masc proteins previously confirmed to be functionally conserved [17, 21, 22, 24, 25] and EkMasc (Fig 6). Due to the high level of homology between the two Masc sequences in *E. kuehniella*, we excluded EkMascB from the analysis, however, a comparison of EkMasc and EkMascB is shown in S9 Fig. The analysis identified two regions with an increased proline content in all species. The first proline-rich domain (PRD-1) is localized between the second zinc finger domain and the masculinizing domain, and the second proline-rich domain (PRD-2) at the C-terminus of the proteins (Fig 6). The *BmMasc* mutant of *B. mori*, which lacked the first 294 amino acids of the protein, was able to masculinize BmN-4 cells, while the same mutant additionally missing the final 75 amino acids failed to do so [19]. As indicated in Fig 7A, this region of the protein corresponds to PRD-2. In addition to PRD-2, we also identified two consecutive tyrosine-asparagine (YN) amino acid residues in this region that are conserved in all six species (Fig 7B). Only in *P. interpunctella* Masc, this tyrosine-asparagine motif is missing, but it should be noted that this region of the *PiMasc* transcript has not yet been confirmed by RT-PCR and the PiMasc protein has not been functionally confirmed.

**Fig 6.**
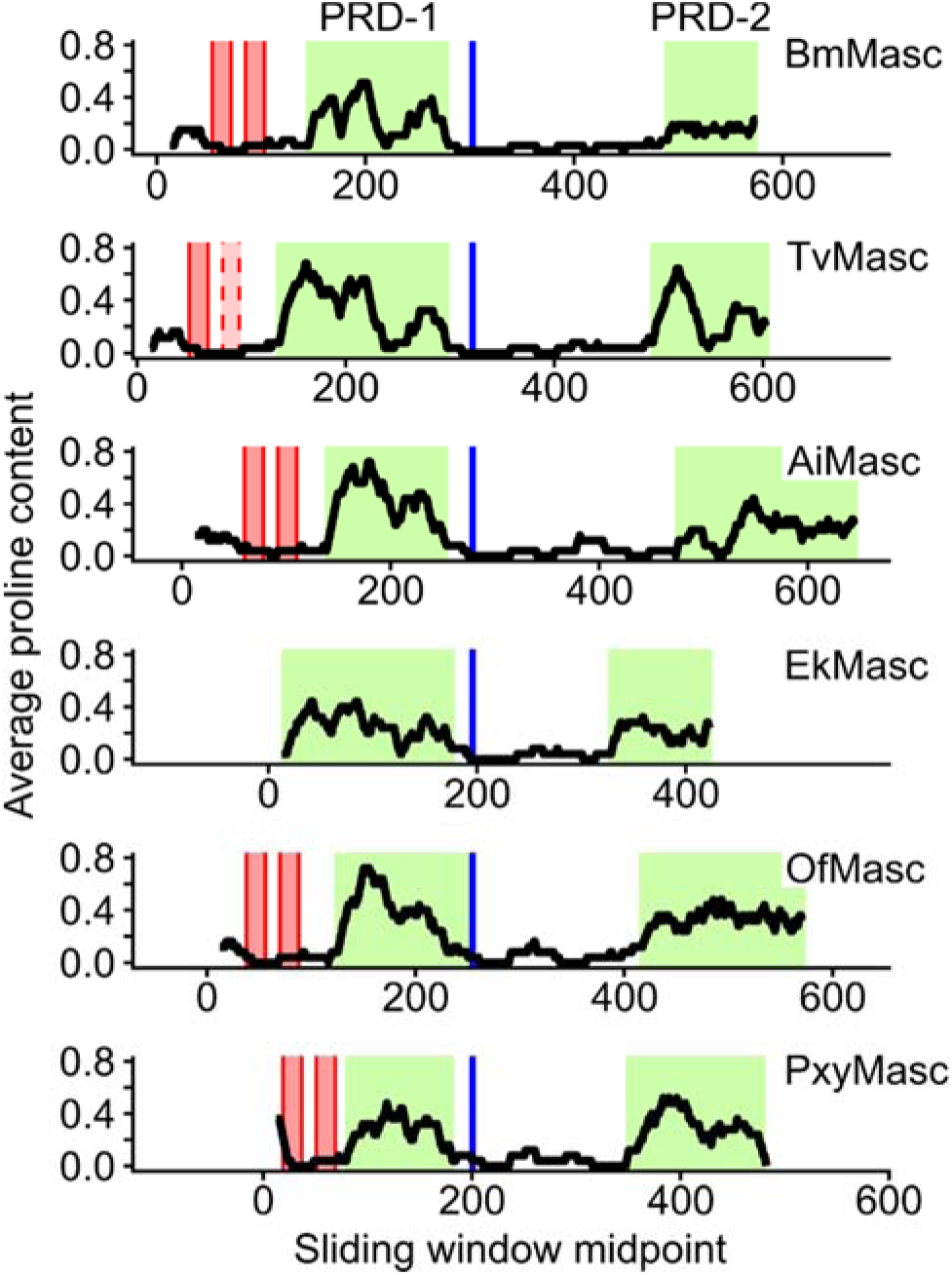
Sliding window analysis of proline distribution in functionally confirmed Masc proteins of six lepidopteran species. Sliding window analysis revealed two regions in all Masc proteins with increased proline content, proline-rich domain 1 (PRD-1) and proline-rich domain 2 (PRD-2) (shaded green). Graphs are aligned by the masculinizing domain (blue). Also indicated are the two zinc finger domains (shaded red with solid outline), the second zinc finger domain in TvMasc has a dashed outline due to a deletion potentially resulting in the loss of the domain. Species used for this analysis: *Bombyx mori* (Bm), *Trilocha varians* (Tv), *Agrotis ipsilon* (Ai), *Ephestia kuehniella* (Ek), *Ostrinia furnacalis* (Of), and *Plutella xylostella* (Pxy).

**Fig 7.**
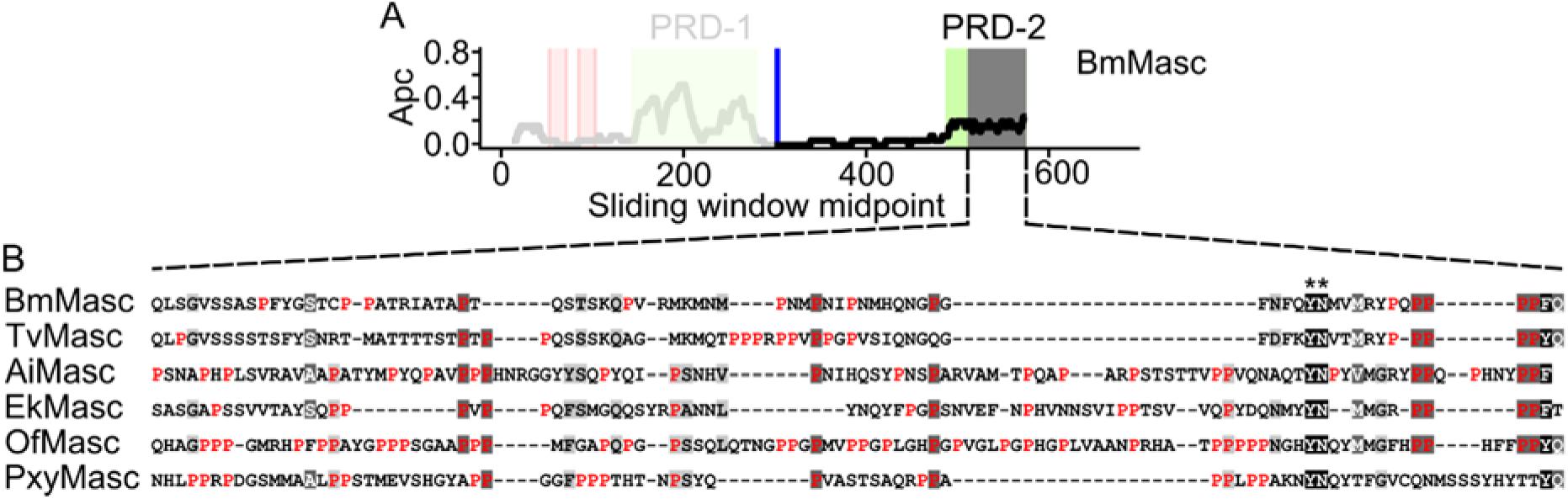
Analysis of the terminal region essential for BmMasc function as identified in Katsuma et al. (2015). In addition to the masculinizing domain (blue), the shaded grey region in (**A**) shows a region essential for Masc protein function, coinciding with the proline-rich domain 2 (PRD-2) (green). This region of BmMasc was aligned to the functionally confirmed Masc protein sequences in other Lepidoptera (**B**). Apart from a tyrosine-asparagine motif (**), which is conserved in all species, the only conserved feature of this region is the relatively high proline (red) content. APC, average proline content. Species used for this analysis: *Bombyx mori* (Bm), *Trilocha varians* (Tv), *Agrotis ipsilon* (Ai), *Ephestia kuehniella* (Ek), *Ostrinia furnacalis* (Of), and *Plutella xylostella* (Pxy).

## Discussion

Duplications of sex determining genes have been described in several insect species. In particular, duplications of the *transformer* (*tra*) gene, the main splicing regulator of *dsx*, are common in the order Hymenoptera [6, 10–12]. In Lepidoptera, only a few species have been investigated, but apart from the primary signal gene in the silkworm, i.e. *Feminizer* (*Fem*), which consists of a high copy tandem repeat generating *Fem* piRNA [17], no duplications of lepidopteran sex determination genes have been reported. Here, we report a duplication of the male-determining gene *Masculinizer* (*Masc*) in *E. kuehniella*. Both the *EkMasc* and *EkMascB* genes lack the tandem zinc finger motif, but do contain the masculinizing domain. We developed a PCR-based genetic sexing method in *E. kuehniella* and analyzed the effects of simultaneous knock-down of these genes on sexual development. Functional analysis showed that the simultaneous knock-down of *EkMasc* and *EkMascB* results in female-specific splicing of *Ekdsx* and lethality in males.

The masculinizing function of *Masc* was initially identified in *B. mori* [17] and is conserved in *Trilocha varians*, *Ostrinia furnacalis*, *Agrotis ipsilon*, and *Plutella xylostella* [21, 22, 24, 25]. Similar to the other species, knock-down of *EkMasc* and *EkMascB* in *E. kuehniella* results in female-specific splicing of *Ekdsx*. The high level of protein similarity between EkMasc and EkMascB indicates that the two proteins likely have the same function in male determination. In particular, the high level of conservation between the proteins in the male-determining domain and the C-terminus suggests a conserved function in sex determination as these protein segments are indicated to be essential for Masc function [19].

Simultaneous knock-down of *EkMasc*/*EkMascB* showed female-specific *Ekdsx* splicing in genetic males, but no feminized males were found among adults. However, when using the siRNA targeting the exon containing the masculinizing domain (siMasc_VII), we observed a highly female-biased sex ratio that is likely due to the male-killing effect caused by the loss of dosage compensation, similar to what was found in *B. mori* [17, 20] and hypothesized in *O. furnacalis* [22] and *P. xylostella* [25]. Surprisingly, this male-killing effect was not observed when using the siRNA targeting exon 2, even though both siRNAs showed similar effects on the *EkMasc*/*EkMascB* expression levels and *Ekdsx* splicing pattern. Whether this is caused by the different exons being targeted by the two siRNAs or due to differences in stability of the siRNAs is currently unknown.

The presence of female-specific *Ekdsx* in genetic males injected with siMasc_II, but the absence of a male-killing effect and lack of persistent phenotypic feminization provide insights into the sex determination mechanism in *E. kuehniella*. Under normal conditions, splicing of *Ekdsx* in ZZ embryos transitions from female-to male-specific at 16–18 hpo, indicating that sex is determined at this point of development. After injections with siMasc_II, splicing of *Ekdsx* was assessed at 48 hpo, showing predominantly female-specific splicing in genetic males 30–32 hours after the sex of the individual is normally established. However, RNAi is transient and the expression levels of *EkMasc*/*EkMascB* likely increased after a number of hours, resulting in a reversal of female-specific *Ekdsx* splicing to male-specific splicing of *Ekdsx >*48 hpo and the development of a male phenotype. This suggests a strong plasticity in timing of sexual development in *E. kuehniella*.

The pattern of sex-specific expression of *Masc* was previously shown only in *B. mori* [17]. In early male embryos, expression of *Masc* increases and subsequently decreases rapidly, while expression in female embryos gradually decreases after oviposition [17]. Here we show that differential expression of *EkMasc* and *EkMascB* also occurs during early embryogenesis in *E. kuehniella*. As found in *B. mori*, the window of differential expression is narrow in *E. kuehniella*, starting at approximately 12–14 hpo, with a maximum around 16 hpo, and ending before 24 hpo. The peak of *EkMasc* and *EkMascB* expression in males corresponds to the first occurrence of the male-specific splice form of *Ekdsx*, suggesting a close interaction between these genes. However, previous studies have shown that dosage maintenance of duplicated genes occurs at both the transcriptional and translational levels in eukaryotes and that repression of translation occurs rapidly rather than gradually [42]. Therefore, it should be noted that even though both *EkMasc* and *EkMascB* show sex-specific differential expression, translational repression might prevent one of the genes to be truly functional. Future research should therefore focus on the separate knock-down/-out of *EkMasc* and *EkMascB* to determine whether one or both genes are involved in sex determination and/or dosage compensation in *E*. *kuehniella*.

We identified a male-specific splice variant of *EkMasc* and *EkMascB* lacking the exon essential for masculinization in *E. kuehniella*, i.e. *EkMasc^ms^* and *EkMascB^ms^*. We further confirmed this splice variant in the closely related *P. interpunctella* (Pyralidae) and the distant codling moth, *C. pomonella* (Tortricidae). In all three species, *Masc^ms^* results from an early stop codon and therefore leads to a truncated protein. Interestingly, an alternative splice variant of *BmMasc* has been identified [43] that lacks the exon containing the *Fem* piRNA target sequence and is therefore not sensitive to the down-regulation by *Fem* piRNA. In *B. mori*, this splice variant is present in both sexes and is essential for the development of the external female genitalia. In contrast, the *Masc^ms^* splice variant is found predominantly in male somatic tissues, but its function is unclear. However, the pattern of splicing and its apparent conservation across Lepidoptera is remarkable. Only in *P. interpunctella* and *C. pomonella*, the Masc^ms^ protein does contain the two zinc finger domains, and therefore in these species Masc might have a tertiary function aside from masculinization and dosage compensation.

The flour moth *E. kuehniella* is the first species described in which the Masc zinc finger motifs were lost naturally. Initially, the dosage compensation function of *Masc* in *B. mori* was predicted to involve the zinc finger domains in the N-terminus of the protein due to their known ability to bind DNA and RNA [17]. However, recent data revealed that the masculinizing domain of BmMasc, rather than the zinc finger domains, regulates dosage compensation [20, 26]. Male-specific lethality of *EkMasc*/*EkMascB*-silenced individuals suggests that dosage compensation also occurs during embryogenesis in *E. kuehniella* and that *EkMasc* and/or *EkMascB* regulate(s) this process either directly or indirectly. This supports the dispensable role of the zinc finger domains in both sex determination and dosage compensation as found in *B. mori* [19, 20, 26]. High conservation in other lepidopteran species, however, suggests an important and conserved function of the zinc finger domains for the Masc protein function.

What this function is, remains currently unknown, but one proposed hypothesis states that the domains increase the binding efficiency of Masc to target RNA or DNA [20]. In addition, they hypothesized that the absence of the zinc finger domain in a mutant strain of *B. mori* might be compensated by another zinc finger-containing gene, *Bmznf*-*2*. Overexpression of *Bmznf*-*2* in BmN cells, derived from female ovarian cells, resulted in partial male-specific splicing of *Bmdsx*, but only if both zinc finger domains were intact [44]. In *E. kuehniella*, we were able to identify a potential ortholog, *Ekznf*-*2*, containing both zinc finger domains (S10 Fig). Therefore, it is possible that the loss of the zinc finger domains in *EkMasc* and *EkMascB* is compensated by the presence of *Ekznf-2* or another zinc finger domain-containing protein. However, why this loss of the zinc finger domains occurred only in *E. kuehniella*, even though *znf-2* is likely to be conserved in Lepidoptera, is unclear. Alternatively, it is possible that the combined expression of *EkMasc* and *EkMascB* can compensate for the potentially reduced efficiency of Masc protein in *E. kuehniella*, thereby negating the loss of the zinc finger domains. Individual knock-out experiments of *EkMasc* and *EkMascB* could provide more clarity regarding the role of the zinc finger domains in *E. kuehniella* and Lepidoptera in general.

Similar to the *transformer* gene in Diptera, Coleoptera, and Hymenoptera, the protein sequence of Masc is highly variable in Lepidoptera. Apart from the previously identified masculinizing domain, the two zinc finger motifs and the bNLS domain, a shared feature of the Masc proteins is the relatively high level of proline content concentrated in two proline-rich domains, PRD-1 and PRD-2. Proline-rich repeats are known to have protein-protein binding capabilities and are often involved in binding of transcription factors [45, 46]. The presence of a proline-rich domain at the C-terminus of the protein is another similarity between Masc and TRA [3], the latter being a confirmed splicing regulator of *dsx* in three insect orders. In addition, it has been shown that not only the masculinizing domain but also a segment of the C-terminus (within the final 75 amino acids) of the protein is essential for masculinization [19]). The C-terminus of Masc coincides with PRD-2, not only in *B. mori* but in all lepidopteran Masc proteins identified. Therefore, we propose that PRD-2, rather than the zinc fingers, of the Masc proteins in conjunction with the masculinizing domain are important for transcriptional regulation and sex determination in Lepidoptera, either through providing protein stability as previously suggested [19] or through binding of transcription factors, or both.

In conclusion, we identified the first duplication of *Masc* in Lepidoptera, *EkMasc* and its paralog *EkMascB*, on the Z chromosome of *E. kuehniella*. Knock-down of *EkMasc* and *EkMascB* revealed male lethality similar to that previously found in other lepidopteran species, confirming that *Masc* is a good target gene to eliminate males, a trait that might be exploited in the future to shift the sex ratio of populations of *E. kuehniella* to increase egg production. In addition, we have identified another, seemingly conserved splice form of *Masc*, *Masc^ms^*, which reveals an additional, previously unknown level of complexity to the lepidopteran sex determination pathway. Overall, our data contribute to the understanding of *Masc* function in *E. kuehniella* and Lepidoptera in general and provide further information regarding *Masc* as a potential target for lepidopteran pest management.

## Materials and methods

### Insects

A laboratory strain (WT-C02) of the Mediterranean flour moth, *E. kuehniella*, was used for all experiments. This wild-type strain was established in 2002 from individuals collected in Boršov nad Vltavou, Czech Republic, and has been maintained on artificial diet [47] under 12:12 (L:D) conditions at 20–22°C. The codling moth, *Cydia pomonella*, samples used were obtained from the laboratory strain Krym-61; its origin and rearing were described earlier [48]. Larvae of the Indian meal moth, *Plodia interpunctella*, were obtained from a local population collected in České Budějovice, Czech Republic.

### Identification and isolation of *EkMasc* and *EkMascB* sequences

To identify *Masculinizer* in *E. kuehniella*, reverse transcription PCR (RT-PCR) was done using RNA from approximately 50 pooled eggs 24 hours post oviposition (hpo) and single male pupa samples, both in duplicate. Total RNA was isolated using TRI Reagent (Sigma-Aldrich, St. Louis, MO) according to the manufacturer’s protocol, using chloroform for phase separation. RNAs were dissolved in 40 μL nuclease-free water and their concentrations were measured on a NanoDrop 2000 spectrophotometer (Thermo Fisher Scientific, Waltham, MA). Approximately 1 μg RNA per sample was converted to cDNA using the ImProm-II Reverse Transcription System kit (Promega, Madison, WI) using the Oligo(dT)15 primer supplied with the kit and with a final concentration of 3 mM MgCl2 according to the manufacturer’s instructions. Primers were designed based on the putative *Masc* transcript sequence of *P. interpunctella* (annotated as maker-scaffold92-augustus-gene-0.127-mRNA-1) obtained through LepBase (lepbase.org; [49]) using Geneious 9.1.6 (https://www.geneious.com; [50]) with default settings. All primers used in the article can be found in S3 Table. PCR was carried out in a final volume of 10 µL containing 0.2 µM of each primer (Masc_F1 and Masc_R1), 0.2 mM of each dNTP, 1× Ex *Taq* PCR buffer, 0.025 units of Ex *Taq* DNA polymerase (TaKaRa, Otsu, Japan), and 1 μL of cDNA. PCR amplification was performed using a standard thermocycling program of 94°C for 3 min initial denaturation; 35 cycles of denaturation at 94°C for 30 s, annealing at 60°C for 30 s, and extension at 72°C for 1 min; with a final extension at 72°C for 3 min. PCR amplification was confirmed by gel electrophoresis using a 1.5% agarose gel in 1× TAE buffer and visualized using ethidium bromide (EtBr). The remaining PCR products were pooled, purified using the Wizard® SV Gel and PCR Clean-Up System (Promega), cloned into the pGEM-T Easy vector (Promega), and finally sequenced (SEQme, Dobříš, Czech Republic) to obtain the initial EkMasc and EkMascB sequences (see S1 Methods for details).

Full-length cDNA sequences of *EkMasc* and *EkMascB* were obtained by rapid amplification of cDNA ends PCR (RACE-PCR) for both 3’- and 5’- ends (see S1 Methods for details). In short, to obtain the 3’ region of the cDNA, first strand cDNA synthesis was prepared using 1 μg of the RNA sample from pooled 24 hpo eggs, the adapter primer (AP), and the ImProm-II Reverse Transcription System (Promega) as described above. Two rounds of PCR amplification followed. In the first round, the Masc_F1 primer and the abridged universal amplification primer (AUAP) were used. In the second round, the primers were substituted by Masc_VII_F1 and AUAP and the template DNA consisted of a 100× diluted PCR product of the first round. For the isolation of the 5’-UTR of *EkMasc* and *EkMascB*, first strand cDNA was synthesized using 1 μg of the same RNA sample with the ImProm-II Reverse Transcription System as described above, but using the Masc_V_R1 primer. The 5’-RACE-PCR was performed as described previously [51]. Full-length cDNA sequences of *EkMasc* and *EkMascB*, two splice variants each, are available (GenBank accession numbers MW505939-MW505942). In addition, all splice variants of *EkMasc* and *EkMascB* were aligned and can be accessed at https://easy.dans.knaw.nl/ui/datasets/… (submitted).

### Genome assembly

To assess the splicing structure and genome organization of *EkMasc* and *EkMascB*, we sequenced the male genome of *E. kuehniella* using Oxford Nanopore technology. DNA was extracted from 5 male larvae using cetyltrimethylammonium bromide (CTAB) DNA extraction as described elsewhere [52] and sequenced on the Nanopore PromethION by Novogene (HK) Co., Ltd. (Hong Kong, China). The sequencing yielded 16.2 Gb of data equivalent to ca. 36.8× genome coverage (assuming a genome size of 440.1 Mb, as determined earlier [53]) with N50 of 24.8 kb. The long reads were deposited in the Sequence Read Archive under accession number PRJNA683200. The reads were assembled using Canu version 1.8 [54] with the following parameters, genomeSize = 440.1m and minReadLength = 10000. The complete cDNA sequences of *EkMasc* and *EkMascB* obtained through RACE PCRs were used in a BLASTn search against the assembled genome and exons were annotated using Geneious 9.1.6. A fragment of the scaffold containing *EkMasc*, *EkMascB*, and an additional 20 kb flanking sequence on either side (sequence submitted to GenBank) was further analyzed using BLASTn in LepBase and NCBI to identify genes and transposable elements. For LepBase, BLASTn searches were only restricted to the CDS database of *P. interpunctella*, while the search using NCBI was restricted to the nucleotide collection of Lepidoptera.

### Assessment of copy number and localization of *Masc* using Southern hybridization

DNA was extracted from single female and male pupae using CTAB DNA extraction as described elsewhere [52]. Concentrations were measured on a Qubit 3.0 Fluorometer using the dsDNA BR Assay Kit (Invitrogen, Carlsbad, CA), and approximately 4 µg DNA per sample was used for DNA digestion reactions. DNA was double digested using *Nde*I × *Not*I, *Dra*I × *Nhe*I (all Fermentas, Vilnius, Lithuania), or *Age*I × *Bsp*HI (New England Biolabs, Ipswich, MA) enzymes (see S1 Methods for details). All digested DNA was separated by electrophoresis on a 1% TBE agarose gel and subsequently transferred by capillary transfer to an Amersham Hybond-N+ nylon membrane (GE Healthcare, Milwaukee, WI).

A probe specific to *EkMasc* with high homology to *EkMascB* (96%) was made by PCR-labeling (see S1 Methods for details). The probe was labeled with digoxigenin-11-dUTPs (Roche Diagnostics, Mannheim, Germany) using primers Masc_Sb_F and Masc_Sb_R. For Southern hybridization, 100 ng of the probe was used. The Southern blot assay was performed as described previously [55] with some modifications [56].

### Z-linkage of *EkMasc* and *EkMascB* by quantitative real-time PCR

To assess the localization of *EkMasc* and *EkMascB* on the Z chromosome, we performed quantitative real-time PCR (qPCR) using genomic DNA as a template and *Acetylcholinesterase 2* (*Ace-2*) as an autosomal reference gene as described previously [39] (see S1 Methods for details). This method relies on the comparison of female and male samples as the ratio between Z-linked genes and autosomal genes is 1:2 in females and 2:2 in males. DNA was isolated from single female and male *E. kuehniella* larvae in triplicate using the NucleoSpin DNA Insect kit (Macherey-Nagel, Düren, Germany) as described elsewhere [57]. The 10 µL qPCR mixture consisted of 1× Xceed SG qPCR Mix Lo ROX (Institute of Applied Biotechnologies, Prague, Czech Republic), 400 nM of each forward and reverse primer (for *EkMasc/EkMascB* primers Masc_F_VIa and Masc_R_VI; for *EkAce-2* primers Ek_Ace2_F and Ek_Ace2_R), and 10 ng of genomic DNA. Data was analyzed as described previously [39]. Briefly, the ratios between the target gene and the autosomal reference gene were calculated using the formula *R* = [(1+*E*Reference)^CtReference^]/[(1+*E*Target)^CtTarget^], where *E* is the primer efficiency and Ct the cycle threshold value. Two hypotheses were tested statistically by an unpaired two-tailed *t*-test for unequal variances: (i) the *EkMasc* and *EkMascB* genes are located on the Z chromosome (the expected female to male ratio is 0.5) and (ii) both genes are autosomal (the female to male ratio is 1). The male and female *R* values were either compared directly, or female values were multiplied by 2 to test both hypotheses. The average *R* value for females and males was calculated and plotted using R version 3.5.2 [58].

### Tissue-specific splicing of *Masc*

After identification of a splice variant encoding a truncated Masc protein, which lacks the two cysteine residues essential for masculinization, i.e. skipping exon VII, we tested the presence of this variant during different developmental stages in *E. kuehniella* females and males. This splice variant is referred to as masculinizing “domain” skipping *Masc* (*Masc^ms^*). For the embryonic stage, we used sexed embryos 16 hpo isolated for the *EkMasc* and *EkMascB* expression analysis (see below). In addition, RNA was isolated from single final instar larvae and single 2- or 3-day-old pupae using TRI Reagent. RNA was subsequently DNase-treated with the Invitrogen TURBO DNA-*free* Kit (Thermo Fisher) according to the manufacturer’s instructions, and 1 μg RNA was converted to cDNA using the ImProm-II Reverse Transcription System as described above. To test splice variation, we performed PCR using primers Masc_bmd_qF1 and Masc_R_X, and 1 μl of cDNA. The same PCR mix and profile were used as for the initial isolation of *EkMasc* and *EkMascB*. PCR products were visualized on a 1.5% TAE agarose gel stained with EtBr.

In addition, we tested whether the *Masc^ms^* splice variant was present in the testis and/or the rest of the body of *E. kuehniella*, *P. interpunctella* and *C. pomonella* (see S1 Methods for identification of *Masc* in *C. pomonella*, primer design, and testes dissection). RNA was extracted and processed as described in the previous paragraph. The obtained cDNA samples were tested by PCR using primers qPiMasc_F1 x qPiMasc_R2 and CpMasc_F4 x CpMasc_R2 for *P. interpucntella* and *C. pomonella*, respectively. Samples of *E. kuehniella* were tested by PCR using primers Masc_bmd_qF1 x Masc_R_X. The PCR mix and thermocycling settings were the same as for the initial identification of *EkMasc* and *EkmascB*, and the PCR products were separated and visualized as described above. The non-sex-specific splice variant of *PiMasc* was uploaded to Genbank (accession number MW505946). In addition, all obtained sequences of *PiMasc* and *CpMasc* were aligned and can be accessed at https://easy.dans.knaw.nl/ui/datasets/… (submitted).

### PCR-based genetic sexing of *E. kuehniella*

Sexing of *E. kuehniella* can be done phenotypically in fifth instar larvae, pupae, and adults, however, sexing during embryogenesis and early larval stages is not possible. Therefore, we developed a PCR-based genetic sexing method for *E. kuehniella.* For this, DNA from three female and three male fifth instar larvae was isolated individually using the NucleoSpin DNA Insect kit (Macherey-Nagel). We observed that the primers Masc_F1 and Masc_R1 targeting *EkMasc* and *EkMascB* consistently showed off-target amplification in female samples, resulting in two bands in female samples and a single, expected band in male samples. We tested these primers at different annealing temperatures and found consistent results at temperatures ranging from 52–60°C with optimal results at 55°C. The 10 µL PCR mix contained 0.2 µM of each primer, 0.2 mM dNTPs, 1× One*Taq* Quick-Load reaction buffer, 0.025 units of One*Taq* DNA polymerase (New England Biolabs), and 5 ng of DNA. The thermocycling profile consisted of 94°C for 3 min initial denaturation; 35 cycles of denaturation at 94°C for 30 s, annealing at 55°C for 30 s and extension at 68°C for 45 s; with a final extension at 68°C for 3 min. This protocol was additionally confirmed using DNA samples from at least 20 adults, 10 pupae and 10 larvae of the WT-C02 strain in multiple independent experiments.

### Expression analysis of *EkMasc* and *EkMascB* during embryogenesis

To measure the expression patterns of *EkMasc* and *EkMascB*, DNA and RNA was simultaneously isolated from single embryos at two-hour intervals, starting at 12 hpo and ending at 24 hpo. RNA was isolated using TRI Reagent according the manufacturer’s protocol and pellets were stored in ethanol at –80°C until further use (see S1 Methods). DNA was simultaneously isolated from the organic phase using the back extraction protocol as described by Thermo Fisher Scientific (see S1 Methods for details). This DNA was used to determine the sex of the individual by PCR using sex-specific markers as described above. After sex determination by PCR, ethanol was removed from the corresponding RNA samples of three to six females and males, and RNA pellets were dissolved in 10 µL of diethyl pyrocarbonate (DEPC)-treated water. The complete RNA samples were DNase-treated with the Invitrogen TURBO DNA-*free* Kit according to the manufacturer’s instructions, and all RNA was used directly for conversion to cDNA using the ImProm-II Reverse Transcription System according to the manufacturer’s instructions. The cDNA was prepared using a combination of Random Primers and Oligo(dT)15 Primer (1:1) supplied with the kit, and with a concentration of 3 mM MgCl2. All cDNA samples were diluted 3× with nuclease-free water before use in qRT-PCR experiments.

qRT-PCR experiments were done on cDNA from the embryo samples using the gene *Ekrp49* as a reference to calculate relative expression levels (see S1 Methods for identification of *Ekrp49* and primer design for qRT-PCR). The experiments were performed as described above for qPCR, but 2 μL of the 3× diluted cDNA was used as a template. A minimum of three biological replicates per sex and time point were used and three technical replicates were used for each sample. Since the number of samples exceeded a single plate experiment, at least five samples (in triplicate) were repeated between plates to correct for between-plate variation, and different genes were measured on separate plates. Relative copy numbers were calculated according to the formula described above in Z-linkage of *EkMasc* and *EkMascB* using qPCR, for *EkMasc* and *EkMascB* separately. An unpaired two-tailed *t*-test with unequal variances was used to test for significant differences in expression levels between sexes of the same age. Statistical analysis and visualization of the data were done using R version 3.5.2 [58].

### Functional analysis of *EkMasc* and *EkMascB*

To assess the function of *EkMasc* and *EkMascB* in sex determination, two short interfering RNAs (siRNAs; see S3 Table) were designed to target *EkMasc* and *EkMascB* simultaneously, based on published recommendations [59, 60]. We tested two siRNAs, one targeting the start of the open reading frame (exon II) and one targeting the exon coding for the two cysteine residues essential for masculinization (exon VII) (see S1 Methods for details on siRNA design). Custom synthetic siRNA duplexes siMasc_II and siMasc_VII (see S3 Table for sequence details) were obtained from Sigma-Aldrich. The siRNA duplexes were dissolved in 100 µL nuclease-free water, then the solution buffer contained 100 mM potassium acetate, 30 mM HEPES, and 2 mM magnesium acetate. In addition, one siRNA published earlier [17], designed against GFP (siGFP), was used as a negative control. All siRNAs were injected individually, no combinations were tested.

Eggs were collected within the first hour after oviposition and injected using a FemtoJet Microinjector (Eppendorf, Hamburg, Germany) (see S1 Methods for details). DNA and RNA were isolated simultaneously from a subset of injected single embryos at 16 hpo (to test knock-down efficiency) or 48 hpo (to test *Ekdsx* splicing, see below) using TRI Reagent as described above in the expression analysis. DNA was used to sex the individuals, and subsequently RNA from a minimum of three confirmed WZ and three confirmed ZZ samples was converted to cDNA. Relative expression levels of *EkMasc* and *EkMascB* were measured and calculated using qRT-PCR as described in the expression analysis. Differences in *EkMasc* and *EkMascB* expression levels between control and knock-down treatments were statistically tested by an unpaired one-tailed *t*-test for unequal variances, for both sexes separately.

Remaining eggs were left to develop and hatched approximately 7-10 days post injection. Larvae were transferred to artificial medium and left to develop until adulthood. The adults were phenotypically sexed and additionally assessed for potential abnormalities in reproductive organs, by dissection using a stereo microscope. In addition, heads of adults were used for CTAB DNA isolation (see above) and genetically sexed as described in section “Expression analysis of *EkMasc* and *EkMascB* during embryogenesis”. The observed adult sex ratios were tested for deviations from the expected sex ratio using the Pearson’s chi-squared test for goodness-of-fit with the expected frequency of 0.5 for both sexes.

To assess the effects of *Masc* knock-down on sex determination, splicing patterns of *doublesex* (*dsx*) are often used [17, 25]. Therefore, we identified a *dsx* ortholog in *E. kuehniella*, (see S1 Methods for details). Primers were designed using Geneious 9.1.6 in exons II and V, flanking the female-specific exons. PCR was done by using the mixture and thermocycling program described above, but with an annealing temperature of 60°C using primers dsx_dR_F2 and dsx_dR_R2 and cDNA from either female or male pupa samples. PCR products, run on a 1.5% TAE agarose gel stained with EtBr, showed alternative splicing between sexes. PCR products of both sexes were purified, cloned, and sequenced as described above. After confirmation of *Ekdsx* by sequencing, we used the same PCR conditions with cDNA from the embryo time series (see “Expression analysis of *EkMasc* and *EkMascB* during embryogenesis”) to identify the timing of sex-specific splicing of *Ekdsx* in *E. kuehniella*. We also used the same PCR on cDNA of a minimum of three WZ and ZZ embryos (of 48 hpo) injected with any of the three siRNAs to determine differential *Ekdsx* splicing. The products were run on a 1.5% TAE agarose gel and stained with EtBr. All obtained *Ekdsx* sequences are included as an alignment and can be accessed at https://easy.dans.knaw.nl/ui/datasets/… (submitted).

### Sliding window analysis of proline content in lepidopteran MASC proteins

We compared all functionally confirmed lepidopteran Masc proteins to identify any other potentially conserved features/domains and we noticed a relatively high proline content in all lepidopteran Masc proteins. To identify patterns in the distribution of these proline residues, we used a sliding window approach. Therefore, we calculated the average proline content across a window of 25 amino acids (aa) and shifted the window by steps of 1 aa until the end of the protein using Microsoft Excel 2013. This process was performed for all functionally confirmed Masc protein sequences, i.e. *Bombyx mori* [17], *Trilocha varians* [21], *Agrotis ipsilon* [24], *Ostrinia furnacalis* [22], and *Plutella xylostella* [25], including EkMasc and EkMascB. An initial trial using window sizes of 10, 20, 25, and 50 aa showed that window size 25 aa was optimal for visualization of the data. The data were visualized using R version 3.5.2 [58] with the ggplot2 package [61], and domains were indicated using Inkscape 0.92 (https://inkscape.org/).

## Supporting information

**S1 Table. Significance test of *EkMasc* and *EkMascB* expression level comparison between sexes during embryogenesis in *Ephestia kuehniella*.** An unpaired two-tailed *t*-test for unequal variances was used. Expression levels significantly differing from each other between sexes (*P* < 0.05) are indicated in bold. (PDF)

**S2 Table. Significance test of knock-down effects of siRNA treatments in *Ephestia kuehniella* using an unpaired one-tailed *t*-test for unequal variances.** Expression levels significantly differing from each other between treatments (*P* < 0.05) are indicated in bold.

(PDF)

**S3 Table. Overview of primers and siRNAs used in this study.**

(PDF)

**S1 Fig. *EkMasc* and *EkMascB* Z-linkage assessed by qPCR using genomic DNA of *Ephestia kuehniella*.** Relative copy numbers were assessed for female and male samples (n = 3 for both sexes). Indicated are hypothetical *EkMasc* and *EkMascB* female to male ratios relative to male copy numbers corresponding to the autosomal hypothesis (both *EkMasc* and *EkMascB* located on autosomes; F:M ratio = 1.00), and the Z chromosomal hypothesis (F:M ratio = 0.50). Error bars indicates standard deviation.

(PDF)

**S2 Fig. Southern blot assay using *EkMasc* probe in *Ephestia kuehniella*.** Two signals can be identified in the genomic DNA of both female and male samples double digested with (1) *Nde*I x *Not*I, (2) *Dra*I x *Nhe*I, and (3) *Age*I x *Bsp*HI. Arrows indicate highly diffused bands. Note that female signals are weaker than male signals. Indicated are the marker (M) in bp, female (♀) and male (♂) samples.

(PDF)

**S3 Fig. Exon-intron map of the *EkMasc* and *EkMascB* genes in *Ephestia kuehniella*.** Two poly-adenylation sites are indicated for each gene, as well as the open reading frame (grey) including the start (ATG) and stop codon (*). Additionally indicated is the splice variant skipping exon VII (dashed grey line) and its corresponding premature stop codon (grey *).

(PDF)

**S4 Fig. Protein alignment of EkMasc and EkMascB of *Ephestia kuehniella*.** Indicated in the alignment are the conserved bipartite nuclear localization signal (bNLS; green) and the masculinizing domain (MD; blue). Also indicated are deletions (red arrows) and insertions (cyan arrows) in either EkMasc or EkMascB based on comparison to the Masc protein sequence of the closely related *Plodia interpunctella*. A single grey arrow (at amino acid 31) indicates an indel between EkMasc and EkMascB that cannot be categorized as a deletion or an insertion in either protein sequence based on the Masc protein sequence of *P. interpunctella*.

(PDF)

**S5 Fig. Genetic sexing of *Ephestia kuehniella*.** Note the two bands in female samples. Indicated are marker (M) in kb, no template control (NTC), female (♀) and male (♂) samples.

(PDF)

**S6 Fig**. **Illustrative set-up of the RNAi experiments in *Ephestia kuehniella*.** Indicated in the figure are the degenerated zinc finger motifs (pink boxes) upstream of the open reading frame, the bipartite nuclear localization signal (green box), the male determining region (blue box), the open reading frame (yellow) and the two siRNAs targeting *EkMasc* and *EkMascB* (red dashed lines). Also shown is an exon representation of the genes.

(PDF)

**S7 Fig. Splicing pattern of *Ekdsx* in *Ephestia kuehniella***. (**A**) Schematic representation of female- (pink line) and male-specific (blue line) splicing patterns of *Ekdsx*. Indicated below the figure are the targets of the primers used throughout the article (F and R). (**B**) The two dominant female- and the dominant male-specific transcripts and their predicted respective open reading frames (in grey/pink). (**C**) Sex-specific splicing of *Ekdsx* as detected by the primers indicated in **A**. A currently uncharacterized third female-specific splice variant is also visible (the top arrow). (**D**) Sex-specific splicing pattern of *Ekdsx* during early development in WZ (left) and ZZ (right) individuals. Note transitions of *Ekdsx* splicing from female-specific to male-specific in ZZ individuals only, 16–18 hours post oviposition (hpo). Indicated are marker (M) in kb, and no template control (NTC).

(PDF)

**S8 Fig. Alignment of functionally confirmed Masc proteins in six lepidopteran species.** The alignment shows low levels of amino acid homology even within the functional domains. Indicated below the alignment are the conserved zinc finger domains (ZF1 & ZF2; pink), the bipartite nuclear localization signal (bNLS; green) and the masculinizing domain (MD; blue). Aligned are the Masc protein sequences of *Bombyx mori* (Bm), *Trilocha varians* (Tv), *Agrotis ipsilon* (Ai), *Ostrinia furnacalis* (Of), *Plutella xylostella* (Pxy), and *Ephestia kuehniella* (Ek). (PDF)

**S9 Fig. Sliding window analysis of proline content for EkMasc and EkMascB proteins in *Ephestia kuehniella*.** Note that the proline distribution in both proteins is very similar.

(PDF)

**S10 Fig. Protein alignment of zinc finger protein 2 (ZNF-2) of *Bombyx mori* (Bm) and the predicted ortholog in *Ephestia kuehniella* (Ek).** Indicated in pink are the two zinc finger domains (ZF1 and ZF2), and in blue two cysteine amino acids separated by two amino acids similar to the masculinizing domain in lepidopteran Masc proteins.

(PDF)

**S1 Methods**

(PDF)

## Acknowledgments

We thank Marie Korchová for excellent technical assistance and Joanna Kotwica-Rolinska for providing technical help with microinjections. We also thank Arjen van’t Hof and Atsuo Yoshido for their valuable comments on the manuscript. Computational resources were supplied by the project “e-Infrastruktura CZ” (e-INFRA LM2018140) provided within the program Projects of Large Research, Development and Innovations Infrastructures.

## Author Contributions

Conceived and designed the experiments: SV, ECV, FM. Performed the experiments and analyzed the data: SV. Assembled the genome of *Ephestia kuehniella*: AV, PN. Funding acquisition: FM. Wrote the paper: SV, ECV, FM.

## Data Availability Statement

The raw Nanopore PromethION long reads from an *Ephestia kuehniella* male generated in this study have been deposited in the Sequence Read Archive under accession number PRJNA683200. All splice variants of the *EkMasc* and *EkMascB* genes and all obtained sequences of the *PiMasc*, *CpMasc*, and *Ekdsx* genes are available at https://easy.dans.knaw.nl/ui/datasets/… (submitted).

## Supporting information

**S1 Table.**
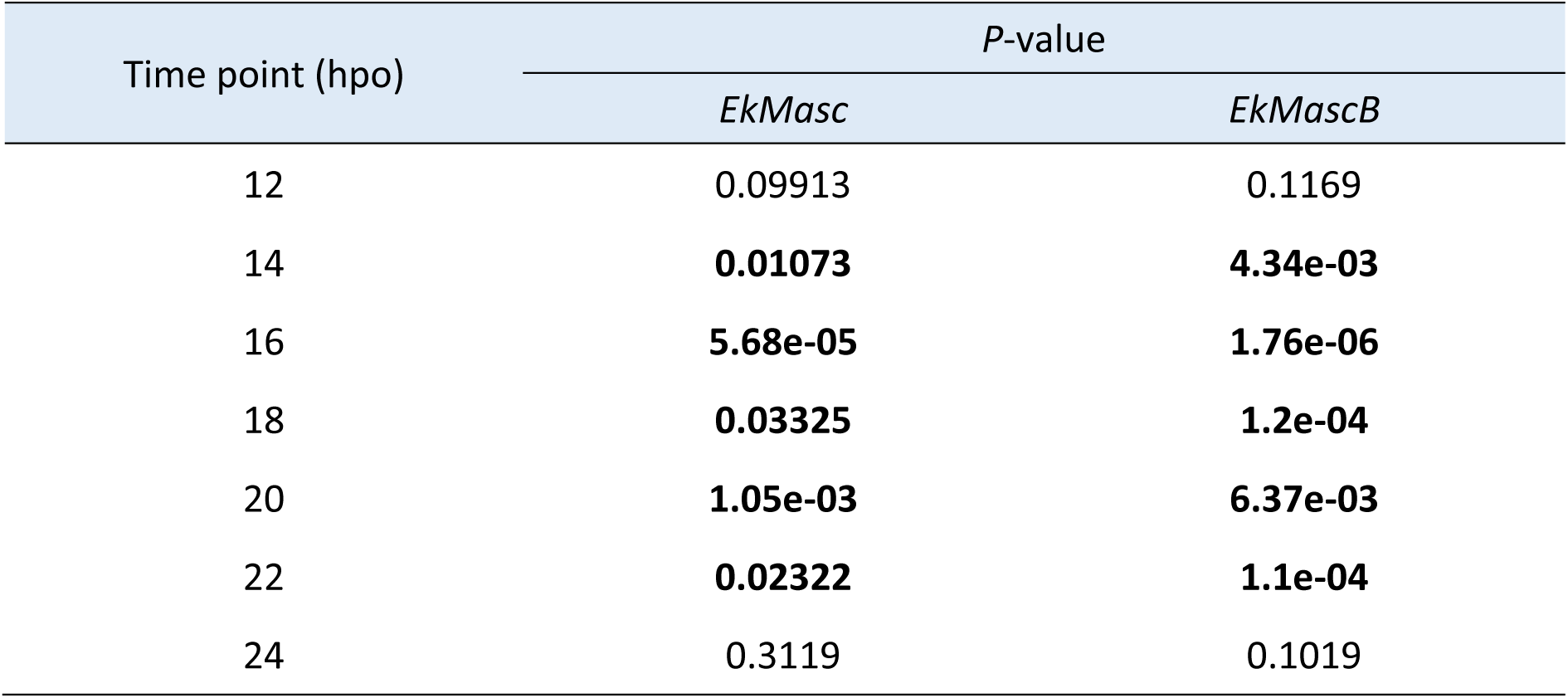
Significance test of *EkMasc* and *EkMascB* expression level comparison between sexes during embryogenesis in *Ephestia kuehniella*. An unpaired two-tailed *t*-test for unequal variances was used. Expression levels significantly differing from each other between sexes (*P* < 0.05) are indicated in bold.

**S2 Table.**
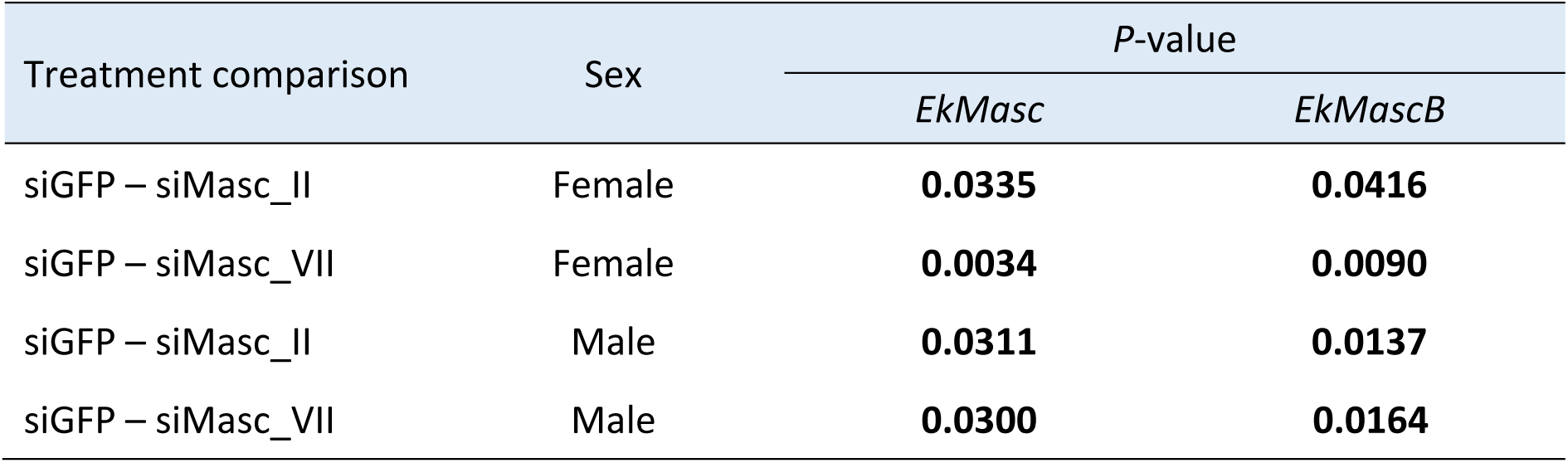
Significance test of knock-down effects of siRNA treatments in *Ephestia kuehniella* using an unpaired one-tailed *t*-test for unequal variances. Expression levels significantly differing from each other between treatments (*P* < 0.05) are indicated in bold.

**S3 Table.**
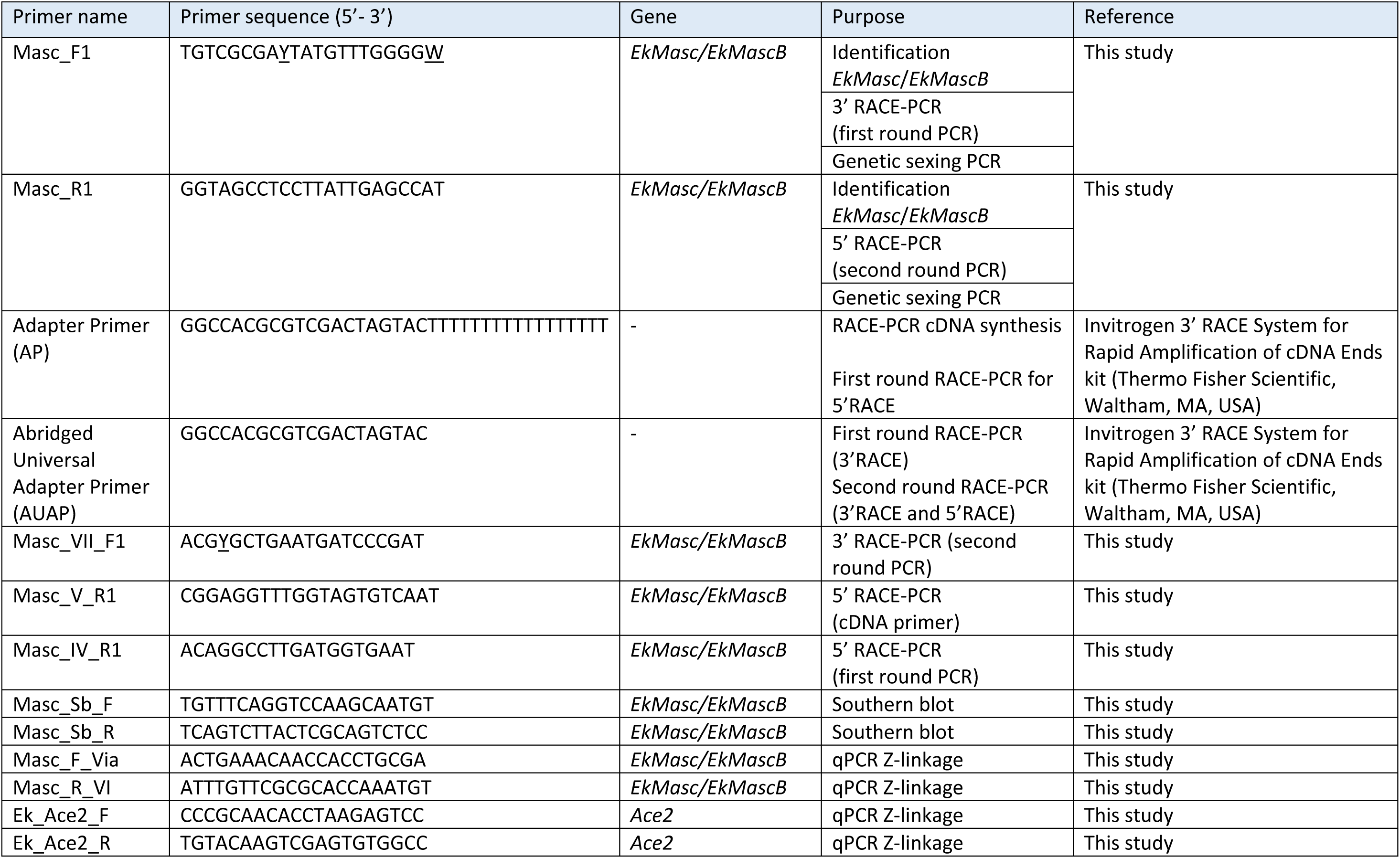
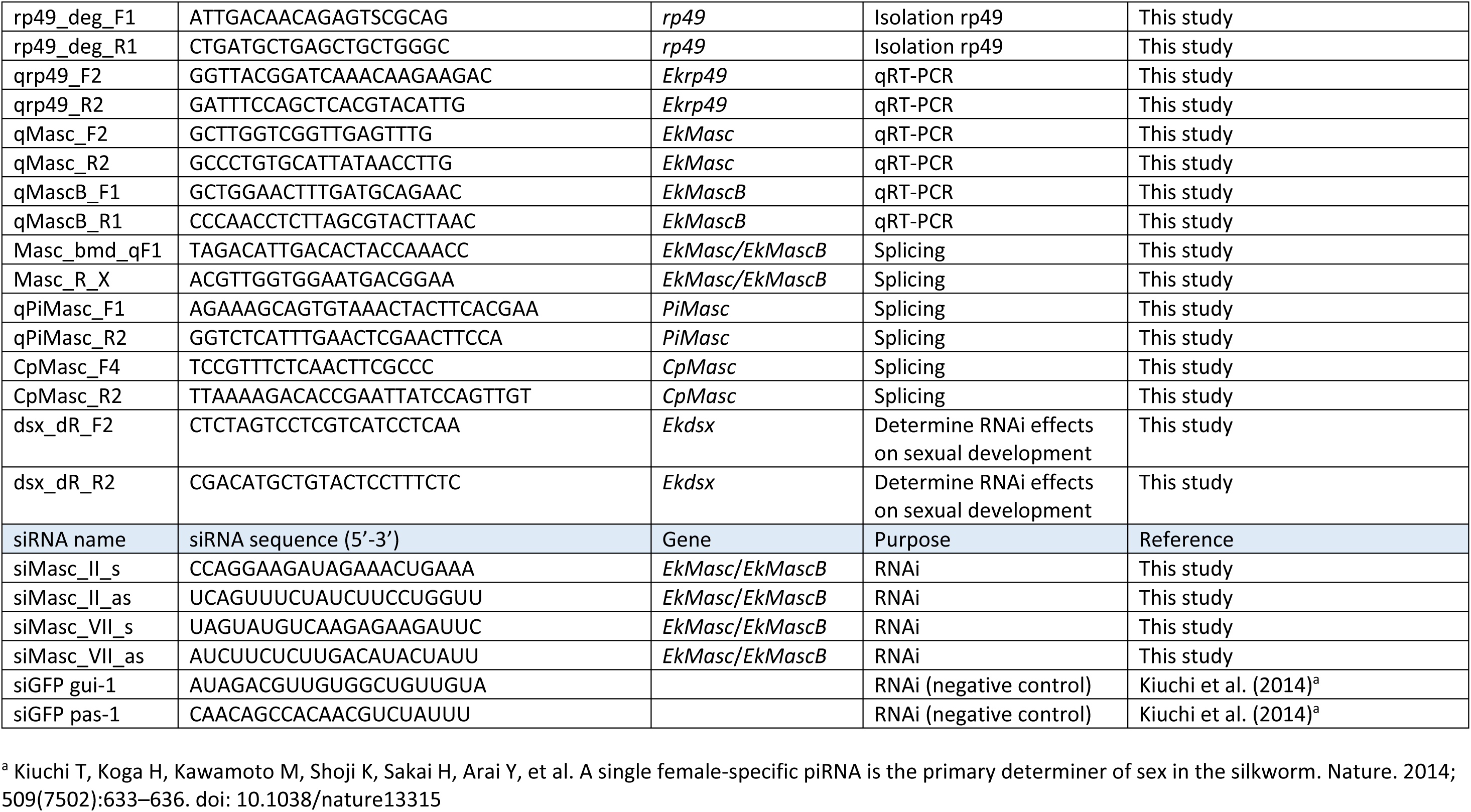
Overview of primers and siRNAs used in this study.

**S1 Fig.**
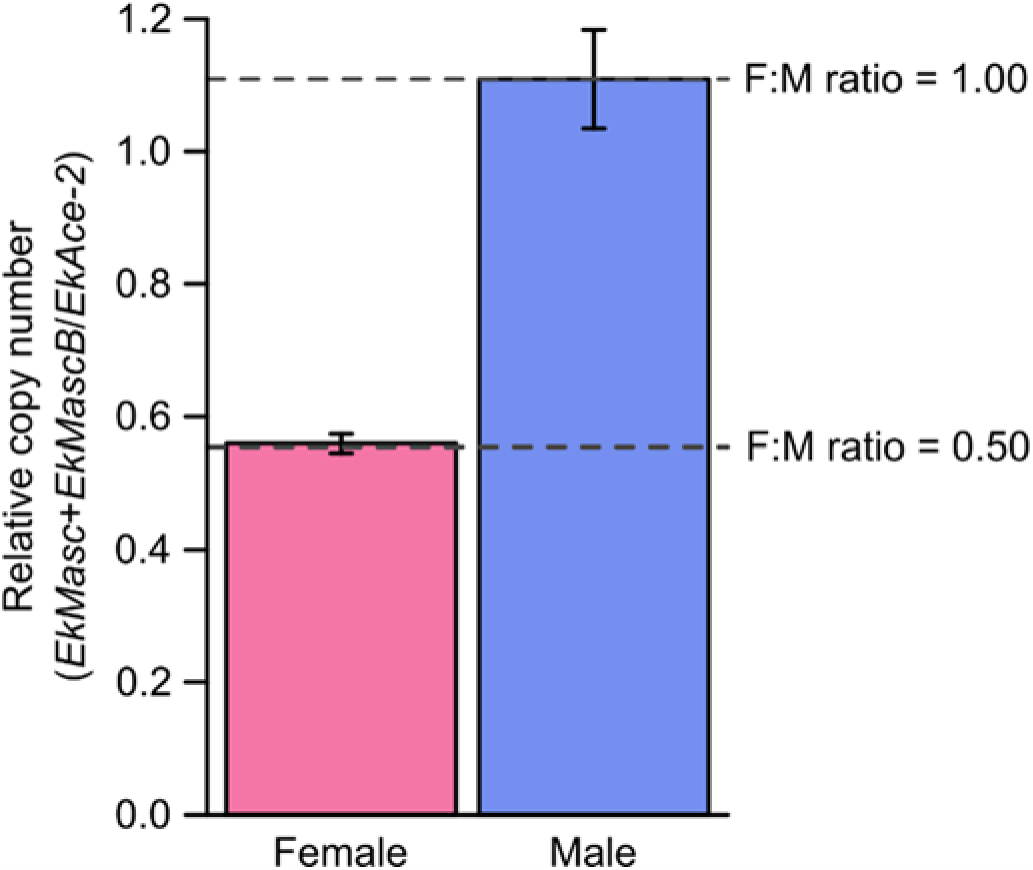
*EkMasc* and *EkMascB* Z-linkage assessed by qPCR using genomic DNA of *Ephestia kuehniella*. Relative copy numbers were assessed for female and male samples (n = 3 for both sexes). Indicated are hypothetical *EkMasc* and *EkMascB* female to male ratios relative to male copy numbers corresponding to the autosomal hypothesis (both *EkMasc* and *EkMascB* located on autosomes; F:M ratio = 1.00), and the Z chromosomal hypothesis (F:M ratio = 0.50). Error bars indicate standard deviation.

**S2 Fig.**
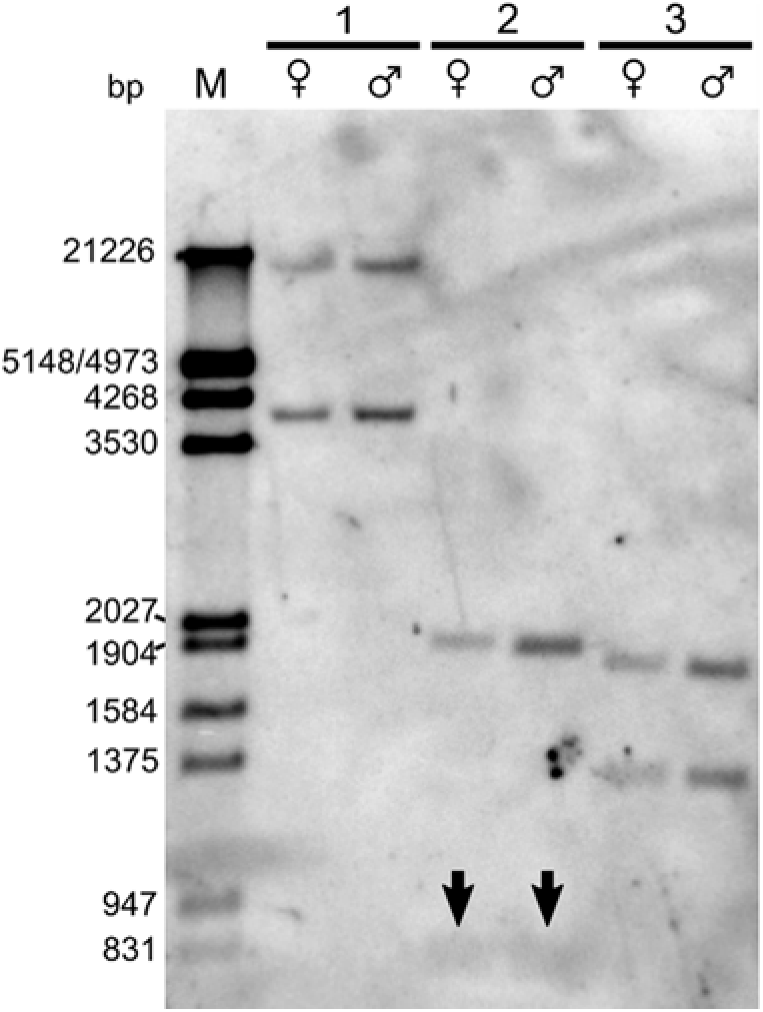
Southern blot assay using *EkMasc* probe in *Ephestia kuehniella*. Two signals can be identified in the genomic DNA of both female and male samples double digested with (1) *Nde*I x *Not*I, (2) *Dra*I x *Nhe*I, and (3) *Age*I x *Bsp*HI. Arrows indicate highly diffused bands. Note that female signals are weaker than male signals. Indicated are the marker (M) in bp, female (♀) and male (♂) samples.

**S3 Fig.**
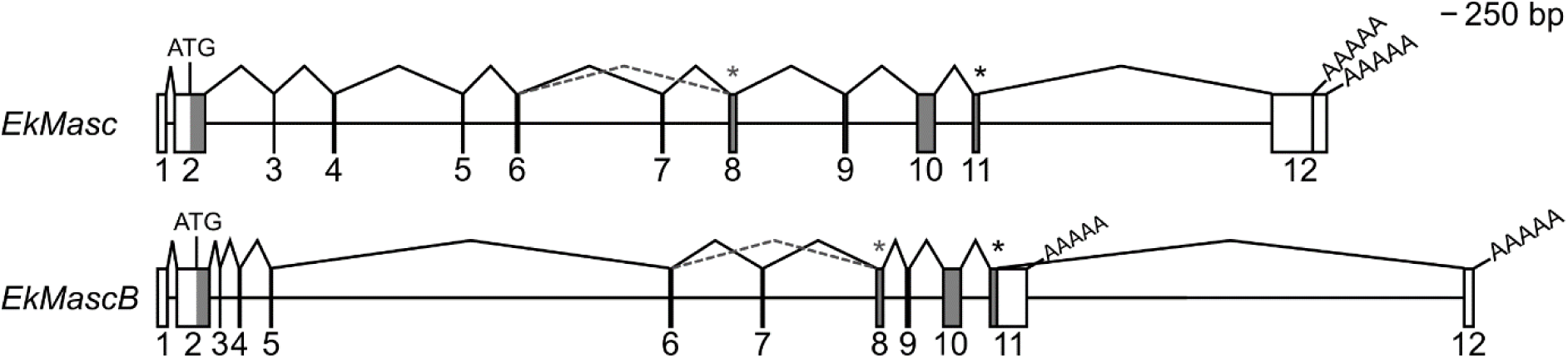
Exon-intron map of the *EkMasc* and *EkMascB* genes in *Ephestia kuehniella*. Two poly-adenylation sites are indicated for each gene, as well as the open reading frame (grey) including the start (ATG) and stop codon (*). Additionally indicated is the splice variant skipping exon VII (dashed grey line) and its corresponding premature stop codon (grey *).

**S4 Fig.**
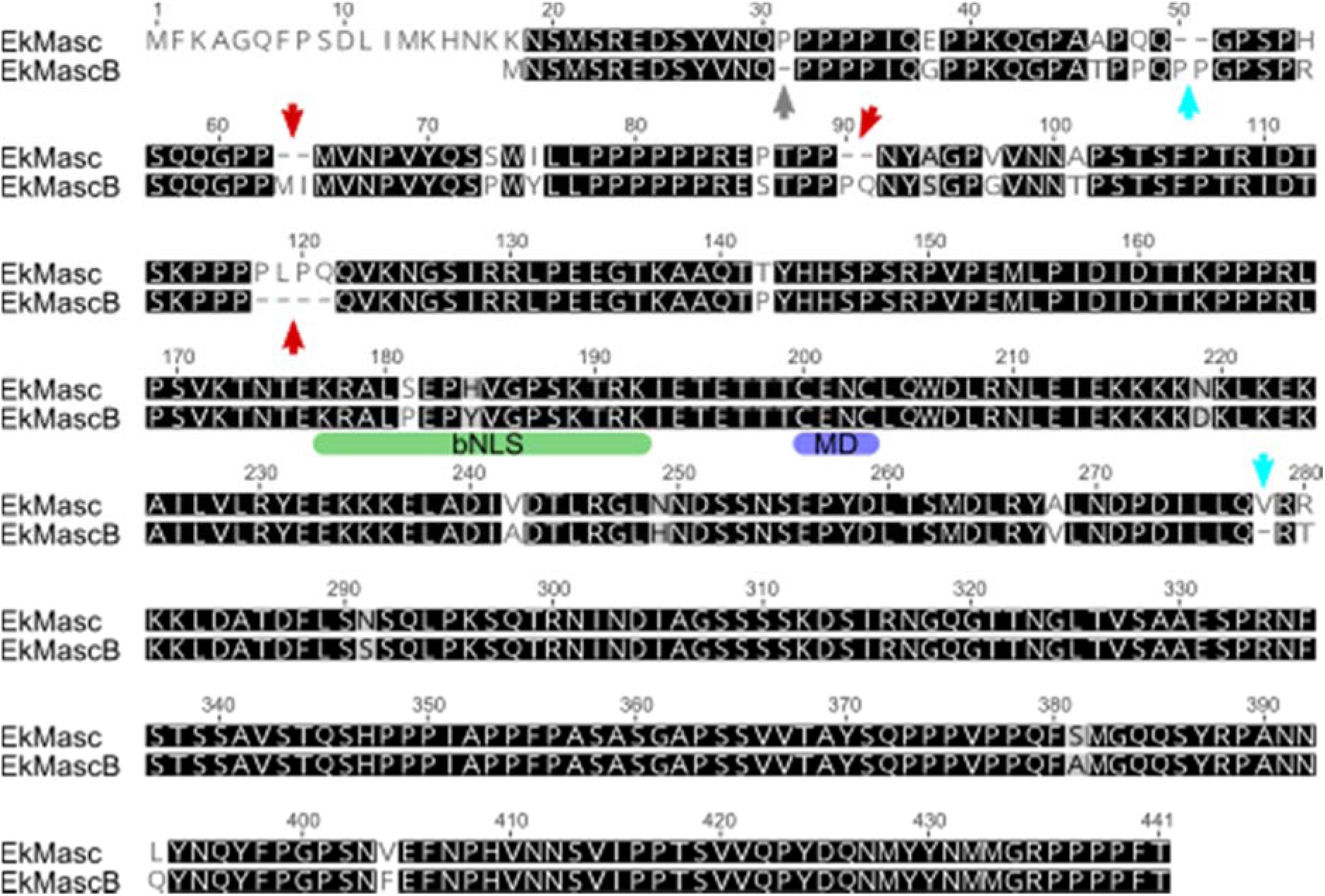
Protein alignment of EkMasc and EkMascB of *Ephestia kuehniella*. Indicated in the alignment are the conserved bipartite nuclear localization signal (bNLS; green) and the masculinizing domain (MD; blue). Also indicated are deletions (red arrows) and insertions (cyan arrows) in either EkMasc or EkMascB based on comparison to the Masc protein sequence of the closely related *Plodia interpunctella*. A single grey arrow (at amino acid 31) indicates an indel between EkMasc and EkMascB that cannot be categorized as a deletion or an insertion in either protein sequence based on the Masc protein sequence of *P. interpunctella*.

**S5 Fig.**
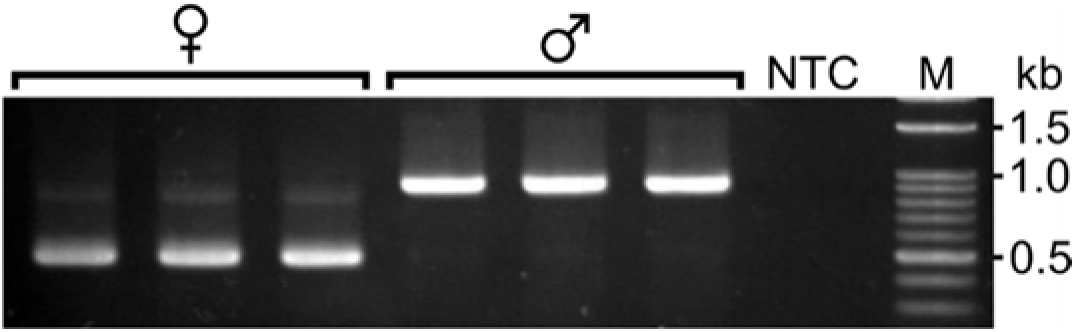
Genetic sexing of *Ephestia kuehniella*. Note the two bands in female samples. Indicated are marker (M) in kb, no template control (NTC), female (♀) and male (♂) samples.

**S6 Fig.**
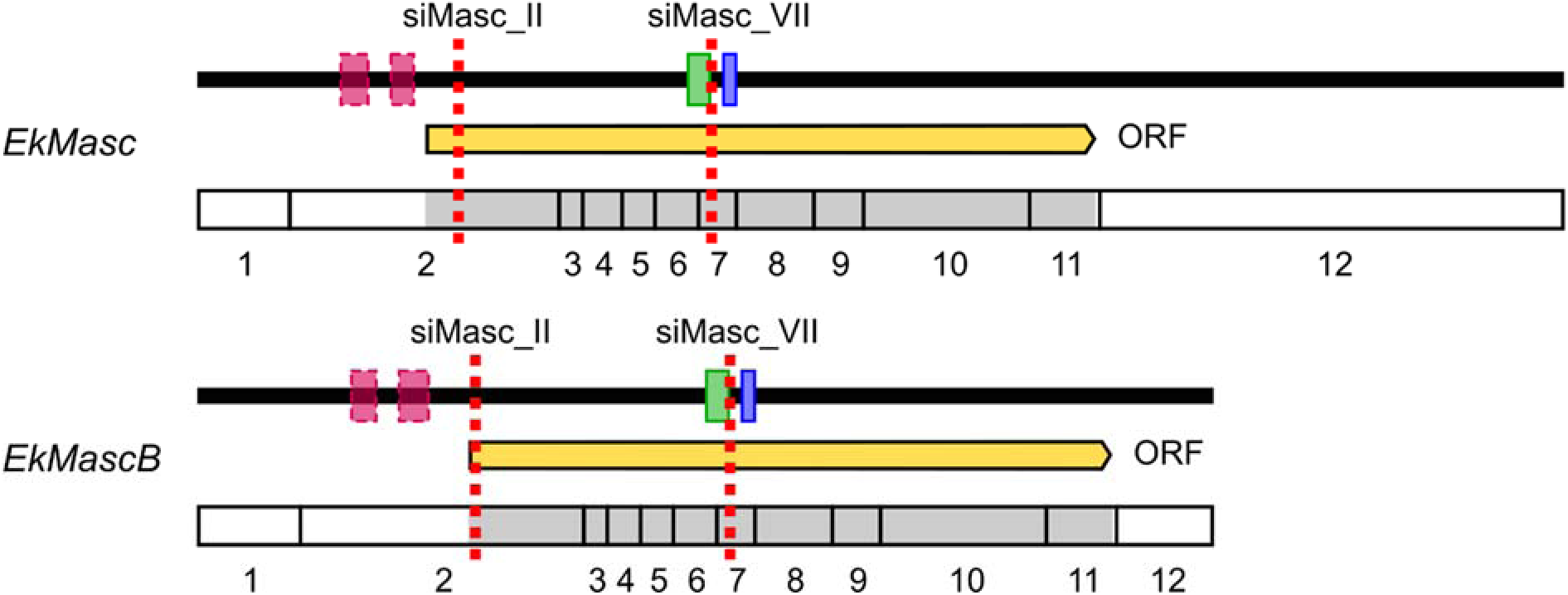
Illustrative set-up of the RNAi experiments in *Ephestia kuehniella*. Indicated in the figure are the degenerated zinc finger motifs (pink boxes) upstream of the open reading frame, the bipartite nuclear localization signal (green box), the male determining region (blue box), the open reading frame (yellow) and the two siRNAs targeting *EkMasc* and *EkMascB* (red dashed lines). Also shown is an exon representation of the genes.

**S7 Fig.**
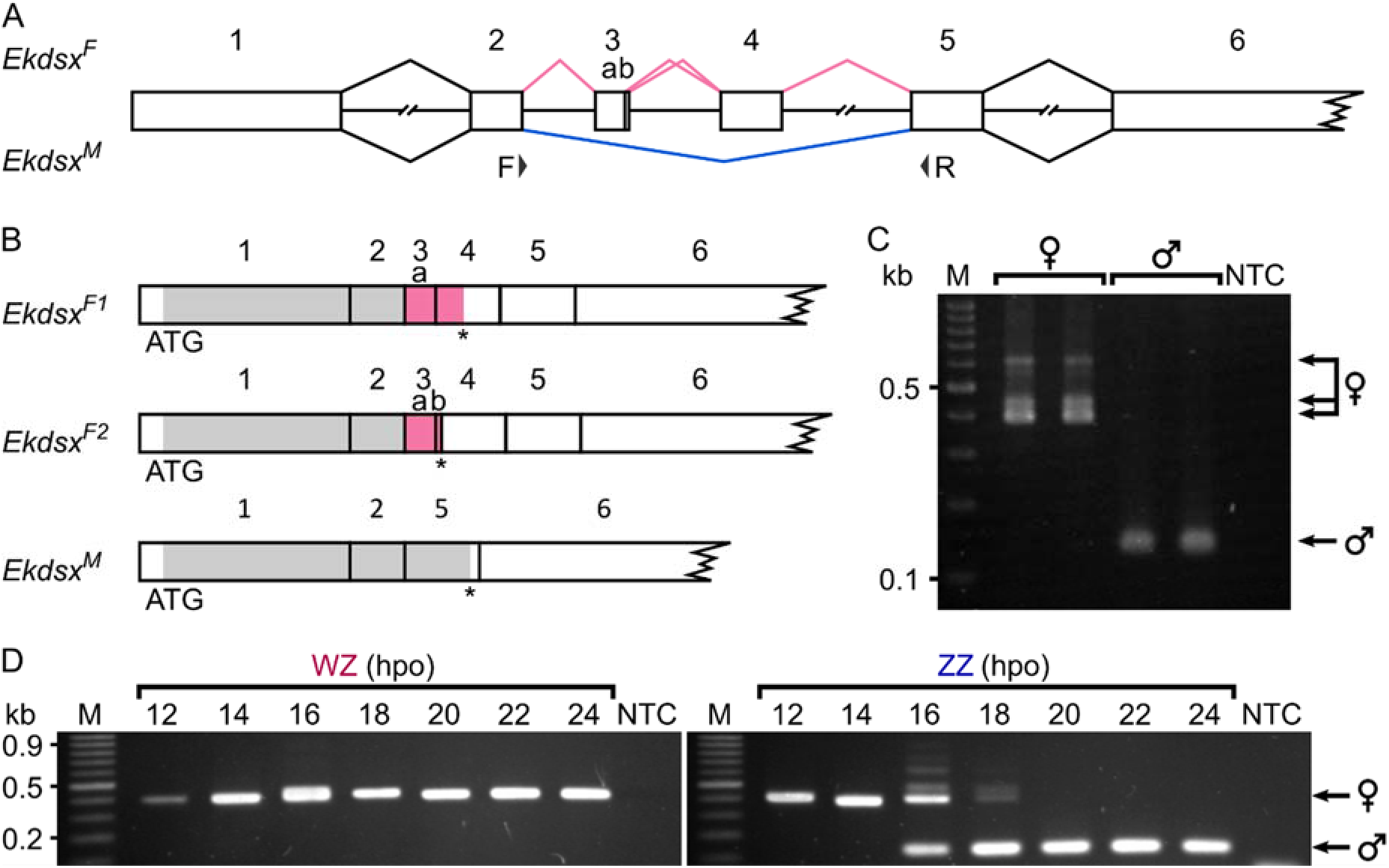
Splicing pattern of *Ekdsx* in *Ephestia kuehniella*. (**A**) Schematic representation of female- (pink line) and male-specific (blue line) splicing patterns of *Ekdsx*. Indicated below the figure are the targets of the primers used throughout the article (F and R). (**B**) The two dominant female- and the dominant male-specific transcripts and their predicted respective open reading frames (in grey/pink). (**C**) Sex-specific splicing of *Ekdsx* as detected by the primers indicated in **A**. A currently uncharacterized third female-specific splice variant is also visible (the top arrow). (**D**) Sex-specific splicing pattern of *Ekdsx* during early development in WZ (left) and ZZ (right) individuals. Note transitions of *Ekdsx* splicing from female-specific to male-specific in ZZ individuals only, 16–18 hours post oviposition (hpo). Indicated are marker (M) in kb, and no template control (NTC).

**S8 Fig.**
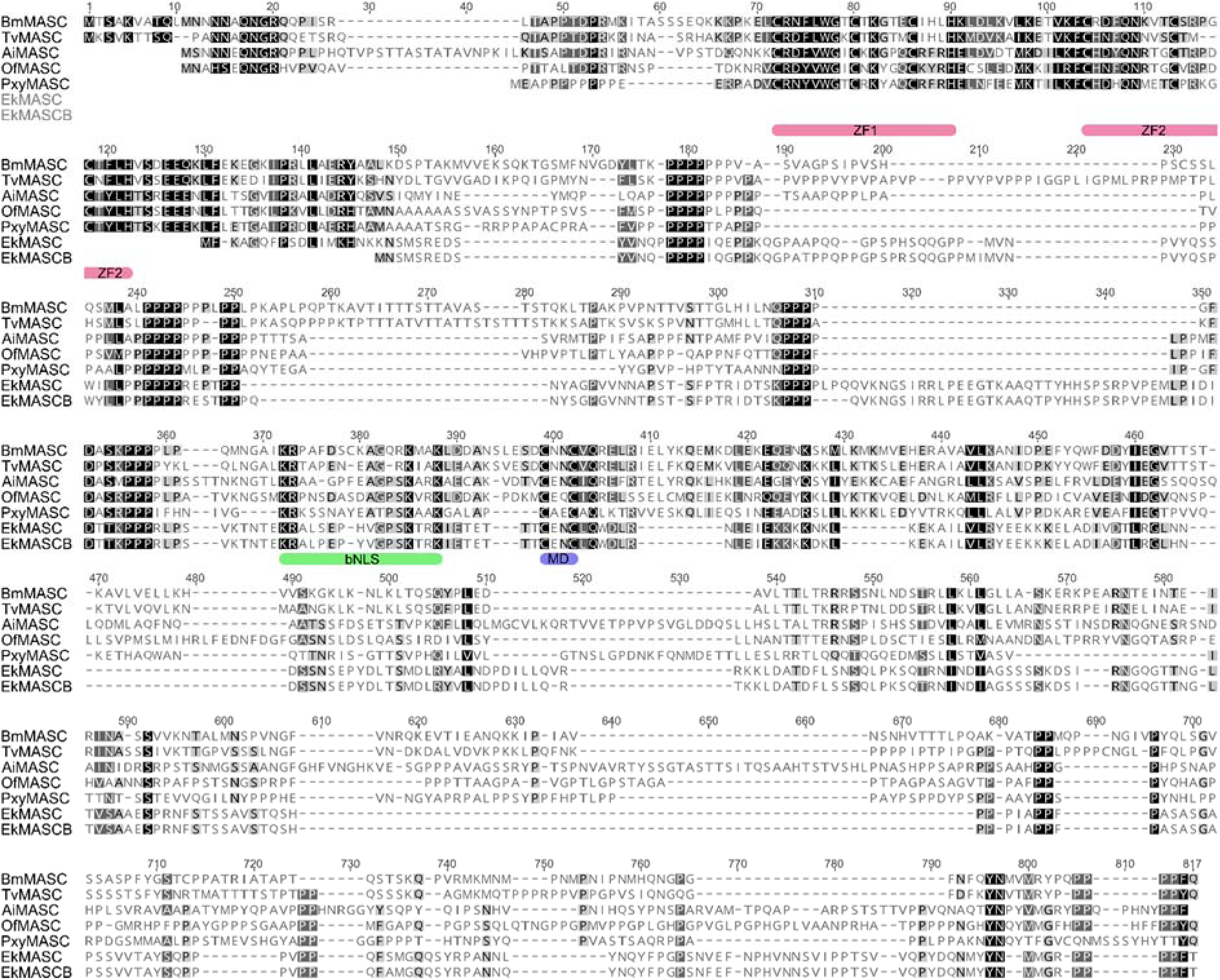
Alignment of functionally confirmed Masc proteins in six lepidopteran species. The alignment shows low levels of amino acid homology even within the functional domains. Indicated below the alignment are the conserved zinc finger domains (ZF1 & ZF2; pink), the bipartite nuclear localization signal (bNLS; green) and the masculinizing domain (MD; blue). Aligned are the Masc protein sequences of *Bombyx mori* (Bm), *Trilocha varians* (Tv), *Agrotis ipsilon* (Ai), *Ostrinia furnacalis* (Of), *Plutella xylostella* (Pxy), and *Ephestia kuehniella* (Ek).

**S9 Fig.**
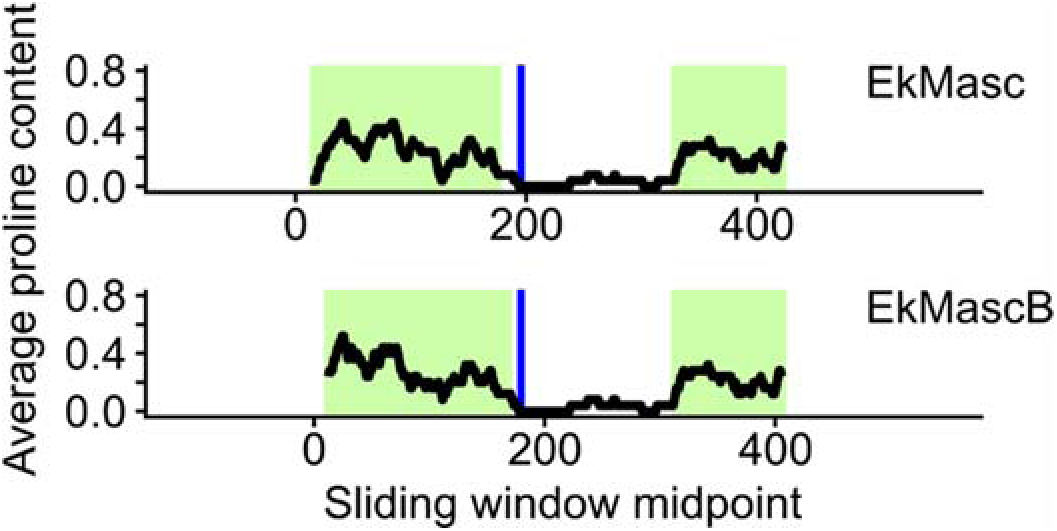
Sliding window analysis of proline content for EkMasc and EkMascB proteins in *Ephestia kuehniella*. Note that the proline distribution in both proteins is very similar.

**S10 Fig.**
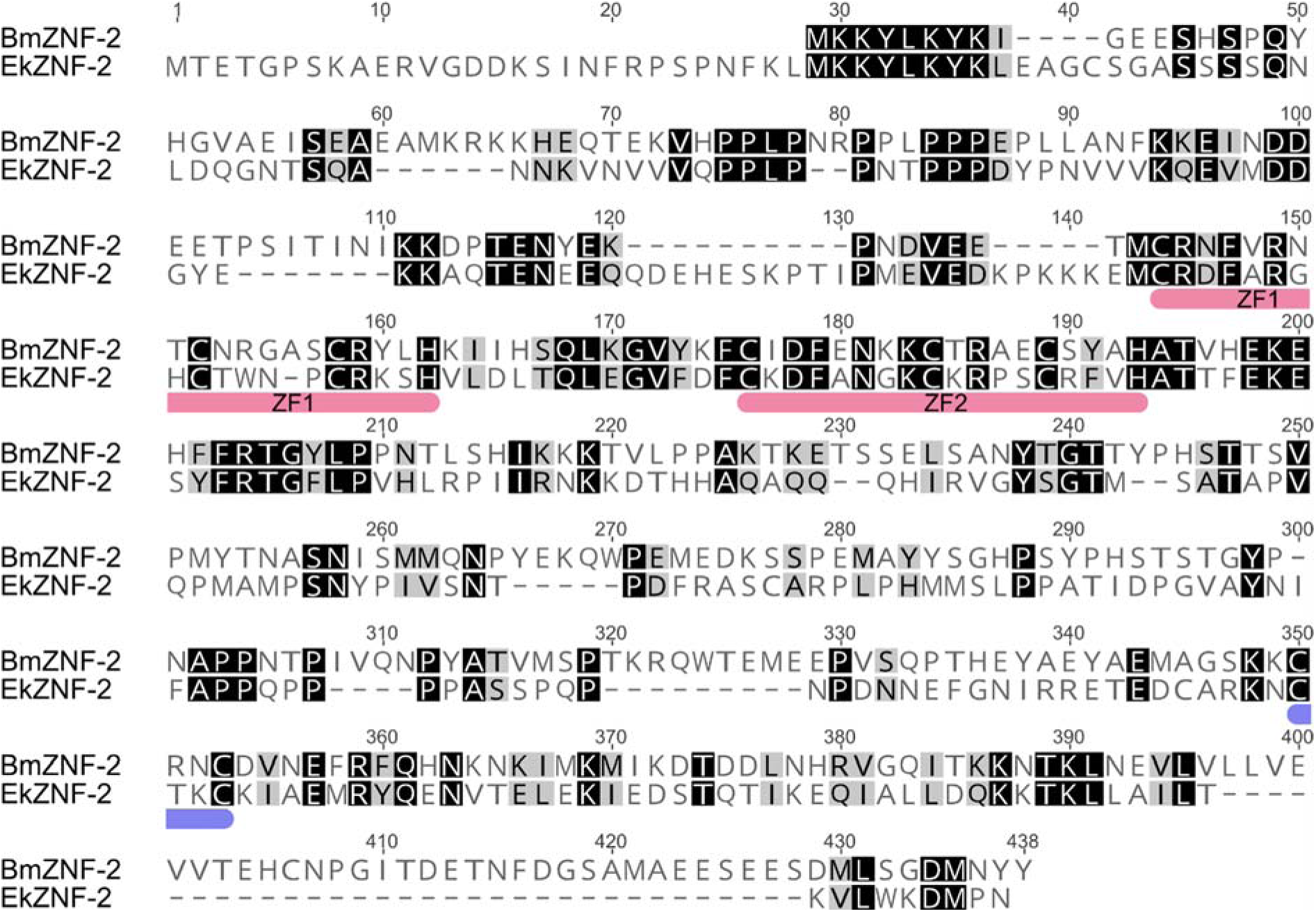
Protein alignment of zinc finger protein 2 (ZNF-2) of *Bombyx mori* (Bm) and the predicted ortholog in *Ephestia kuehniella* (Ek). Indicated in pink are the two zinc finger domains (ZF1 and ZF2), and in blue two cysteine amino acids separated by two amino acids similar to the masculinizing domain in lepidopteran Masc proteins.

## S1 Methods

### Identification and isolation of *EkMasc* and *EkMascB* sequences

#### Cloning, plasmid isolation and sequencing

The PCR products were pooled (eggs and pupa samples separate) and purified using the Wizard® SV Gel and PCR Clean-Up System (Promega, Madison, WI, USA) and subsequently cloned by ligating overnight into the pGEM-T Easy vector (Promega) and transforming into *Escherichia coli* strain DH5α according to the manufacturer’s protocol with minor adjustments. Cells were heat-shocked for 90 s at 42°C, and only 800 μL of LB medium was added during the recovery phase. Positive colonies were screened by PCR using universal M13 primers, and amplicons were checked on a 1% agarose gel. Colonies with inserts of the predicted size were inoculated into 3 mL of LB medium containing 50 μL/mL ampicillin and incubated overnight at 37°C, 200 rpm. Plasmids were subsequently isolated using the NucleoSpin Plasmid kit (Macherey-Nagel, Düren, Germany) according to the manufacturer’s protocol and inserts were sequenced (SEQme, Dobříš, Czech Republic).

#### RACE-PCR

The first round of 3’-RACE-PCR was performed with a gene-specific forward primer (Masc_F1) and the abridged universal amplification primer (AUAP). This 10 µL final PCR reaction mixture was composed of 2 µM gene-specific forward primer, 0.2 µM AUAP, 0.8 mM dNTPs, 1× Ex *Taq* DNA polymerase buffer, 0.25 units of Ex *Taq* DNA polymerase and 1 μL of template cDNA. The primer concentration of the forward primer was increased with respect to the reverse primer to avoid off-target amplification of only the reverse primer. An additional round of semi-nested PCR was performed to increase gene-specific amplification. The PCR reaction volume was increased to 20 µL and the forward primer was substituted with a nested gene-specific primer (Masc_VII_F1) at a final concentration of 0.2 µM (equal to the reverse primer). In addition, the amount of Ex *Taq* DNA polymerase was increased to 0.5 units per reaction. Template for the semi-nested PCR was 100× diluted non-purified PCR product of the preceding 3’ RACE-PCR. Both PCR reactions were performed at 94°C for 3 min; 35 cycles of 94°C for 40 s, 55°C for 40 s and 72°C for 2 min 30 s; and a final extension at 72°C for 3 min. Final PCR products were visualized on a 1% agarose TAE gel using ethidium bromide (EtBr) as a stain. PCR products were purified directly or strong bands were cut from the gel and purified using Wizard® SV Gel and PCR Clean-Up System (Promega) as described by the manufacturer, cloned and sequenced as described above.

For 5’-RACE-PCR, we followed the procedure described in Frohman et al. (1988). The synthesized cDNA was purified using an Illustra Sephadex G-50 column (GE Healthcare Life Sciences, Buckinghamshire, UK) and then poly-A tail was added using Terminal Transferase (New England Biolabs, Ipswich, MA, USA) according to the manufacturer’s protocol with 200 µM dATPs and an incubation time of 45 min. Amplification of gene specific 5’ RACE transcripts was done in two steps. The first step was performed in a 10 µL PCR volume and was composed of 0.2 µM adapter primer (AP), 0.2 µM gene-specific primer Masc_IV_R1, 0.8 mM dNTPs, 1× Ex *Taq* DNA polymerase buffer, 0.25 units of Ex *Taq* DNA polymerase and approximately 10-30 ng of poly-A-tailed cDNA. For the second step, PCR volumes were increased to 20 µL, primers were substituted with AUAP and the nested gene-specific primer Masc_R1, and 1 µL of the 10× diluted PCR-product from the previous step was used as template DNA. Both thermocycling reactions were performed as described above for 3’ RACE PCR. Final PCR products were processed as described for 3’ RACE-PCR.

### Assessment of copy number and localization of *Masc* using Southern hybridization

#### DNA digestion for Southern hybridization

DNA was double digested using *Nde*I × *Not*I, *Dra*I × *Nhe*I (all Fermentas, Vilnius, Lithuania), or *Age*I × *Bsp*HI (New England Biolabs) following the manufacturer’s instructions, using Orange buffer, Tango buffer, or CutSmart buffer, respectively. Digestions were incubated at 37°C overnight and an additional volume of these enzymes was added the next morning, after which the samples were incubated for another 1 h to ensure complete digestion of the DNA. Digestion reactions were stopped by adding Gel Loading Dye Purple (New England Biolabs) to a final 1× concentration. Then the digested DNA was separated on a 1% TBE agarose gel, run at 70 V per centimeter of gel distance for 4 h.

#### Southern hybridization probe

Primers for the Southern hybridization probe were designed in the region of high sequence identity between *EkMasc* and *EkMascB* targeting exon XI using Geneious 9.1.6. Initial PCR was performed using Masc_Sb_F and Masc_Sb_R primers and male genomic DNA as a template using the standard PCR mix described above (see “Identification and isolation of *EkMasc* and *EkMascB* sequences”). The obtained PCR products were checked on a 1% agarose gel for successful amplification, purified, cloned, and sequenced as described above. Plasmid DNA was isolated from a clone containing the expected *EkMasc* sequence and this plasmid DNA was used in PCR to generate a template for PCR-labeling. This 50 µL PCR reaction consisted of 0.2 µM of both Masc_Sb_F and Masc_Sb_R, 0.2 mM dNTPs, 1× Ex *Taq* Buffer, 0.25 units of Ex *Taq* DNA polymerase, and 5 ng of plasmid DNA. The reaction was performed using the thermocycling program described for the initial isolation of *EkMasc* and *EkMascB*. Products were purified using the Wizard® SV Gel and PCR Clean-Up System according the manufacturer’s instructions and used as a template in the labeling reaction. The PCR-labeling reaction consisted of 400 nM each of Masc_Sb_F and Masc_Sb_R, 40 µM each of dATP, dCTP and dGTP, 14.4 µM dTTP, 25.6 µM of digoxigenin-11-dUTPs (Roche Diagnostics, Mannheim, Germany), 1× Ex *Taq* Buffer, 0.625 units of Ex *Taq* polymerase, and approximately 5 ng of template in a total reaction volume of 25 µL. Amplification of the probe was done according to the following profile: denaturation at 94°C for 90 s (1 cycle), denaturation at 94°C for 30 s, annealing at 55°C for 30 s, elongation at 72°C for 60 s (35 cycles), and final elongation at 72°C for 60 s (1 cycle). The probes were then purified using an Illustra Sephadex G-50 column, and their concentrations were measured on a Qubit 3.0 Fluorometer using the dsDNA BR Assay Kit (Invitrogen, Carlsbad, CA).

### Z-linkage of *EkMasc* and *EkMascB* by quantitative real-time PCR (qPCR)

#### qPCR

Concentrations of DNA extracted from *Ephestia kuehniella* larvae were measured on an Invitrogen Qubit 3.0 Fluorometer using the dsDNA BR Assay Kit. Primers were designed to target both *EkMasc* and *EkMascB* simultaneously in a conserved region of the genes using Geneious 9.1.6. The protein sequence of *Bombyx mori* acetylcholinesterase type 2 (accession number ABY50089.1) was used in a tBLASTn search against the *E. kuehniella* genome to identify a partial *EkAce-2* sequence (accession number MW505944), which was then used to design primers using Geneious 9.1.6. qPCR was carried out on a C1000 Thermal cycler CFX96 Real-Time System (Bio-Rad, Hercules, CA) with a cycle program of 95°C for 3 min initial denaturation followed by 45 cycles of 94°C for 30 s denaturation, 60°C for 20 s combined annealing and extension, 95°C for 15 s final denaturation, and 65°C to 95°C by 0.5°C steps of 5 s to analyze the melting curve. Experiments were run in FrameStar 96 well plates sealed with qPCR adhesive foil (both Institute of Applied Biotechnologies, Prague, Czech Republic). Primer efficiencies were determined by dilution series analysis and calculated using the Bio-Rad CFX Manager 3.0 software (Bio-Rad Laboratories, Hercules, CA).

### Tissue-specific splicing of *Masc*

#### Primer design

Primers were designed using Geneious 9.1.6 with “Product Size” setting to 450–500 for *Cydia pomonella* and 1000–1200 for *Plodia interpunctella*. To ensure the design of primers that would amplify both *Masc* and *Masc^ms^* splice variants, the “Target Region” setting was set to include the exon containing the masculinizing region in both species. To this end, the *CpMasc* sequence was manually assembled from the non-assembled transcriptome reads, as the tBLASTn search against the assembled transcriptome did not yield any significant hits. An initial fragment of *CpMasc* was obtained by a tBLASTn search against the non-assembled *C. pomonella* 1-day-old eggs transcriptome reads (accession number SRX5284305), after which the initial sequence was extended using multiple rounds of BLASTn searches until an open reading frame was detected (accession number MW505945).

#### Dissection

We dissected out the testes of the pre-final instar larvae (prior to fusion of the testes) of *E. kuehniella*, *P. interpunctella*, and *C. pomonella* in physiological solution (Glaser 1917; cited in Lockwood, 1961). For each individual, we immediately isolated RNA from the testes and separately from the remaining body tissue using TRI Reagent as described in Materials and methods (see main text). For testis samples, 20 μg of RNA grade glycogen (Thermo Fisher Scientific, Waltham, MA) was added before precipitation to reduce loss of RNA. In *E. kuehniella*, the same procedure was repeated for pupa samples.

Lockwood AP. ‘Ringer” solutions and some notes on the physiological basis of their ionic composition. Comp Biochem Physiol. 1961; 2:241–289. doi: 10.1016/0010-406x(61)90113-x

### Expression analysis of *EkMasc* and *EkMascB* during embryogenesis

#### Sample treatment

For each time point, sixteen embryos were crushed individually in TRI Reagent and stored at –80°C until RNA was isolated to ensure a minimum of three samples for each sex would be available to measure expression levels. Prior to precipitation, 20 µg glycogen RNA grade (Thermo Fisher Scientific) was added as co-precipitate to reduce RNA loss. RNA pellets were stored at –80°C in ethanol until further use.

#### Back extraction DNA isolation protocol

#### https://www.thermofisher.com/cz/en/home/references/protocols/nucleic-acid-purification-and-analysis/dna-extraction-protocols/tri-reagent-dna-protein-isolation-protocol.html

In short, an equal volume of back extraction buffer (4 M guanidine thiocyanate, 50 mM sodium citrate, 1 M Tris base) was added to the organic phase, samples were mixed for 15 s, centrifuged, and the aqueous phase was transferred to a new tube. 20 µg of glycogen was added to the samples and DNA was precipitated by adding 2/3 volume of isopropanol and incubating for 5 min at room temperature. After centrifugation, the pellets were washed twice with 70% ethanol and dissolved in 20 µL nuclease-free water.

#### Identification of *Ekrp49* and primer design

A set of degenerated primers (rp49_deg_F1 and rp49_deg_R1) was designed for *rp49* in a conserved region of the gene in Lepidoptera based on sequences from *B. mori* (NM_001098282.1), *Glyphodes pyloales* (MH715949.1), *Helicoverpa armigera* (JQ744274.1), and *Heliconius melpomene* (EF207973.1). The PCR mixture and cycling program were the same as described for the initial isolation of *EkMasc* and *EkMascB*, but with an annealing temperature of 52°C and scaled up to a final volume of 40 μL. The products obtained were purified, cloned and sequenced as described above. Primers for *Ekrp49* (qrp49_F2 and qrp49_R2) were designed on the *E. kuehniella* sequence obtained (accession number MW505943) using Geneious 9.1.6, while primers for *EkMasc* (qMasc_F2 and qMasc_R2) and *EkMascB* (qMascB_F1 and qMascB_R1) were designed manually in highly diverged regions of the genes. Manually designed primers were subsequently checked for primer-dimers, hairpins, and off-target amplification using Geneious 9.1.6. All primers were tested with single embryo samples and pooled embryo samples for potential off-target amplification and primer-dimers prior to the experiment. To detect off-target amplification and primer-dimers, melting curves were analyzed for secondary peaks and products were run on a 1.5% agarose gel. In addition, amplification efficiencies of each primer pair were determined by a three-fold dilution series of the pooled embryos cDNA sample using the Bio-Rad CFX Manager 3.0 software (Bio-Rad Laboratories).

### Functional analysis of *EkMasc* and *EkMascB*

#### siRNA design

*EkMasc* and *EkMascB* coding sequences were aligned and screened for two consecutive adenine nucleotides, conserved in both sequences, in exons II and VII. For both exons, additional consecutive 19 nucleotides of perfect homology between the two sequences were selected as potential siRNA target sequences and were compared against the *E. kuehniella* genome using BLASTn to identify potential off-target binding sites. For both siRNA duplexes, the lowest homology to any other sequence in the genome (<15 nt perfect homology), GC-content, and sequence asymmetry in the siRNA duplexes were assessed.

#### Egg collection and microinjection

Freshly emerged adult females and males were collected and left to mate overnight. Mating couples were isolated the next morning. Because *E. kuehniella* females lay eggs at dusk, the females were transferred to a Petri dish approximately 5 min before the lights off and eggs were collected by 45 min later. Glass microscope slides were prepared by placing a wet piece of filter paper near each end of the slide, and eggs were aligned against the paper using a slightly wetted paint brush. Injections were done using a FemtoJet Microinjector (Eppendorf, Hamburg, Germany). Needles were prepared from 1 mm outer diameter, 0.58 mm inner diameter borosilicate glass capillaries with filament (Sutter Instrument, Novato, CA, USA) using a Magnetic Glass Microelectrode Horizontal Needle Puller PN-31 (Narishige, Tokyo, Japan), as described in Kotwica-Rolinska et al. (2019). After injection, filter papers were removed, and the glass slides were transferred to a Petri dish containing wetted tissue paper to keep high humidity levels. Petri dishes were incubated at 21–22°C.

Kotwica-Rolinska J, Chodakova L, Chvalova D, Kristofova L, Fenclova I, Provaznik J, et al. CRISPR/Cas9 genome editing Introduction and optimization in the non-model insect *Pyrrhocoris apterus*. Front Physiol. 2019; 10:891. doi: 10.3389/fphys.2019.00891

#### Identification of *Ekdsx*

To identify a *dsx* ortholog in *E. kuehniella*, we used the male DSX protein from *B. mori* (BmDSXM; accession number AHF81625.1) to perform a tBLASTn search against the *E. kuehniella* genome. However, the male-specific protein segment did not provide any significant hits. Therefore, we used BmDSXM to perform a BLASTp search against all lepidopteran species and obtained the predicted DSXM protein of the closely related *Amyelois transitella* (accession number XP_013184257.1). This AtDSXM sequence was subsequently used to identify the male-specific segment of *Ekdsx*.

